# *Leishmania major*-induced alteration of host cellular and systemic copper homeostasis drives the fate of infection

**DOI:** 10.1101/2023.07.25.550461

**Authors:** Rupam Paul, Adrija Chakrabarty, Suman Samanta, Swastika Dey, Raviranjan Pandey, Saptarshi Maji, Aidan T. Pezacki, Christopher J. Chang, Rupak Datta, Arnab Gupta

## Abstract

Copper plays a key role in host-pathogen interaction. We found that during *Leishmania major* infection, the parasite-harboring macrophage regulates its copper homeostasis pathway in a way to facilitate copper-mediated neutralization of the pathogen. Copper-ATPase ATP7A transports copper to amastigote-harboring phagolysosomes to induce stress on parasites. *Leishmania* in order to evade the copper stress, utilizes a variety of manipulative measures to lower the host-induced copper stress. It induces deglycosylation and degradation of host-ATP7A and downregulation of copper importer, CTR1 by cysteine oxidation. Additionally, *Leishmania* induces CTR1 endocytosis that arrests copper uptake. In mouse model of infection, we report an increase in systemic bioavailable copper in infected animals. Heart acts as the major organ for diverting its copper reserves to systemic circulation to fight-off infection by downregulating its CTR1. Our study explores reciprocal mechanism of manipulation of host copper homeostasis pathway by macrophage and *Leishmania* to gain respective advantages in host-pathogen interaction.

## Introduction

Copper (Cu), an essential micronutrient, is vital for any biological system. It is necessary for the functioning of several important enzymes playing critical roles in physiology, e.g., cytochrome c oxidase, dopamine β-hydroxylase, tyrosinase, ceruloplasmin. The copper-dependent enzymes use the capacity of copper to switch between two redox states Cu (II) and Cu (I), to catalyse metabolic processes. Due to its two redox states, the transition between them can lead to a Fenton-like reaction that generates reactive species, resulting in oxidative damage to the cell^1^. Organisms have developed mechanisms whereby certain dedicated proteins are employed to strictly manage the bioavailability of copper. They are equipped with copper transporters, chaperones, and other regulatory proteins that concertedly maintain copper homeostasis^2, 3^.

A cell primarily acquires copper through the high-affinity copper transporter CTR1, which also participates in reducing Cu (II) to Cu (I) to make it bioavailable^4, 5^. The copper is then sequestered by several cellular copper chaperones and transported to designated areas of the cell via several different pathways. ATOX1 is a major cytosolic copper chaperon which transports copper to the two Cu-ATPases located on the Golgi membrane, known as ATP7A (present in all cells except liver) and ATP7B (primarily in liver)^6^. Under normal condition, ATP7A and ATP7B transport copper in the Golgi lumen to the cuproproteins as they mature. However, when copper levels increase in the cell, the Cu-ATPases vesicularize from the Golgi and remove excess copper from the cell via lysosomal exocytosis^7^. Apart from this secretory pathway, copper ion can also bind to the copper chaperone Cox17, which transports it to the mitochondrial cytochrome c oxidase. Additionally, CCS protein transfers Cu to Cu/Zn SOD^6^.

Besides its biosynthetic role, the redox property of copper has been exploited by many mammalian hosts to fight-off a variety of pathogenic infection. We have previously reported that the parasitic kinetoplast, *Leishmania* is subjected to copper stress by its host macrophage in order to neutralize it^8^. *Leishmania* is a digenetic protozoan that causes a wide spectrum of neglected tropical diseases collectively known as leishmaniasis. According to WHO, about 700000 to 1 million new cases occur annually. These numbers occurring across about 100 endemic countries put immaculate pressure on the global healthcare system. Disease symptoms vary from disfiguring skin lesions to life-threatening enlargement of visceral organs, depending on species type^9–11^. *Leishmania* alternates between the female sandfly and mammalian hosts as flagellated promastigotes and intracellular amastigote respectively. *Leishmania* enters mammalian host *via* sand fly bite and are rapidly phagocytosed by macrophages^12^. Phagocytosis is followed by promastigote to non-flagellated amastigote transformation in the phagolysosomal compartments. Amastigotes keep on proliferating within the acidic phagolysosomes until the parasite load causes the cell to burst, thereby, spreading infection^13, 14^. *Leishmania* is known to survive and proliferate inside this compartment by withstanding hostile factors like lysosomal hydrolases, free radicals and low pH^15^. They not only alter their own gene expression but also manipulate the host machinery for their survival^16–18^. The dearth of vaccine and growing resistance to the existing drugs makes it necessary to study and understand the molecular machinery and physiology of *Leishmania*^19, 20^.

Previous research have demonstrated that Cu exerts copper toxicity, which plays a significant role in host-pathogen interaction and the removal of pathogens. Free copper can produce ROS and NO, which can then chemically react with different elements and acids in the cell to magnify the toxic effects of these species^21^. Additionally, it has been observed that Cu-deficient hosts are more vulnerable to both eukaryotic infections like *Candida albicans* and *Trypanosoma lewisi* and prokaryotic pathogens like *Pasteurella hemolytica* and *Salmonella typhimurium*^22–25^. Copper deficiency also impairs the ability of neutrophils and macrophages to neutralise bacteria^26, 27^. Studies reveal that *Mycobacterium avium* and *Escherichia coli* infection of macrophages results in copper channelization to the phagosomal compartment, indicating that hosts are likely to use copper to protect against intracellular pathogens^28, 29^. In a similar vein, mutations in the pathogenic CuATPase in *E. coli* that result in deficiencies in the Cu exporting system increase the vulnerability of the pathogen to Cu-mediated death, highlighting the significance of copper in host-pathogen interactions^29^.

P-type Cu-ATPases can sense increased copper levels and remove them from the cell, making them crucial for maintaining copper homeostasis. The ATPases, CopA and CopB carry out this copper export in bacteria^30, 31^. *N. fowleri*, *Plasmodium falciparum* and *C. albicans* were also shown to use their copper transporter or Cu-ATPase to detoxify copper and maintain homeostasis. In *Saccharomyces cerevisiae*, a single Cu-ATPase known as Ccc2p exists while ATP7 is the sole Cu-ATPase found in *Drosophila*^32^. ATP7A and ATP7B branched out from ATP7 with increasing tissue complexity in higher eukaryotes^33^. Our group has previously characterized the CuATPase in *Leishmania major* and named it *LmATP7* and further identified how it contributes to its survival in host. It is first full-length CuATPase to be functionally characterised in the Trypanosomatid group. The pathogen’s ability to infect and survive in host decreased when *LmATP7* was knocked down. It was further demonstrated that at 24 hours post-infection, host CuATPase, ATP7A traffics from the Golgi, where it normally resides, and localises to the *Leishmania*-positive compartments, possibly to exert Cu stress on the pathogen^8^.

In the present study, we demonstrate that host ATP7A trafficking to pathogen-harboring compartments lead to copper accumulation in them. While dissecting ATP7A response to infection, we observe an overall parasite-induced perturbation in the host copper secretory pathway. ATP7A and primary copper importer CTR1 protein levels were lowered by *Leishmania* via distinct mechanisms in an effort to reduce host-induced copper toxicity. Knock-down experiments confirmed the importance of these proteins in restricting infection. Finally, by using mouse model, we confirmed that copper bioavailability and its proper channelization from various organs to the site of infection are crucial factors to limit *Leishmania* infection.

## Results

### *Leishmania major* infection triggers trafficking of macrophage ATP7A

ATP7A is the primary copper transporting ATPase in macrophages. Previous studies have demonstrated that host macrophage utilizes copper against intracellular pathogens like *Mycobacterium avium* and *Escherichia coli*^28, 29^. Macrophage uses its copper ATPase, ATP7A to neutralize pathogenic yeasts like *Candida* sp^27^. Mammalian ATP7A is a well-characterized copper ATPase that carries out two distinct functions. Under normal physiological conditions, it transports copper to copper-requiring enzymes like lysyl oxidase, dopamine-β-hydroxylase, cytochrome C oxidase, and tyrosinase^34, 35^. When faced with excess intracellular copper, ATP7A vesicularizes out of Golgi to export copper from the cell (Figure S1A). Previously, studies from our group have shown that a fraction of macrophage ATP7A traffics out of Golgi 24 hours post-*Leishmania major* infection^8^. This post-Golgi vesicular movement of ATP7A to *Leishmania*-harboring compartment was attributed to mobilization of copper for killing the pathogen.

In this study, we began by investigating the copper distribution inside *Leishmania-*infected macrophages by using the copper sensor dye, CF4. CF4 binds to Cu and emits at 585 nm. The intensity and distribution of fluorescence is directly proportional to the intracellular copper content and localization respectively^36^. In post-3 hours infected cells, we observed strong, clustered signal (magenta) around intracellular amastigote compartments compared to the widely distributed signal in uninfected macrophages (Figure 1A). ATP7A vesicularization (in green) was observed in the same infected macrophages. The dispersed ATP7A signal is indicative of its copper transport function in the vesicular compartments (Figure 1B). The findings indicate that copper was indeed channelized to the *Leishmania*-positive compartments in infected macrophages. Upon measuring intracellular copper content, we found an increase, though not statistically significant in *Leishmania* infected cells (Figure 1C).

**Figure 1.**
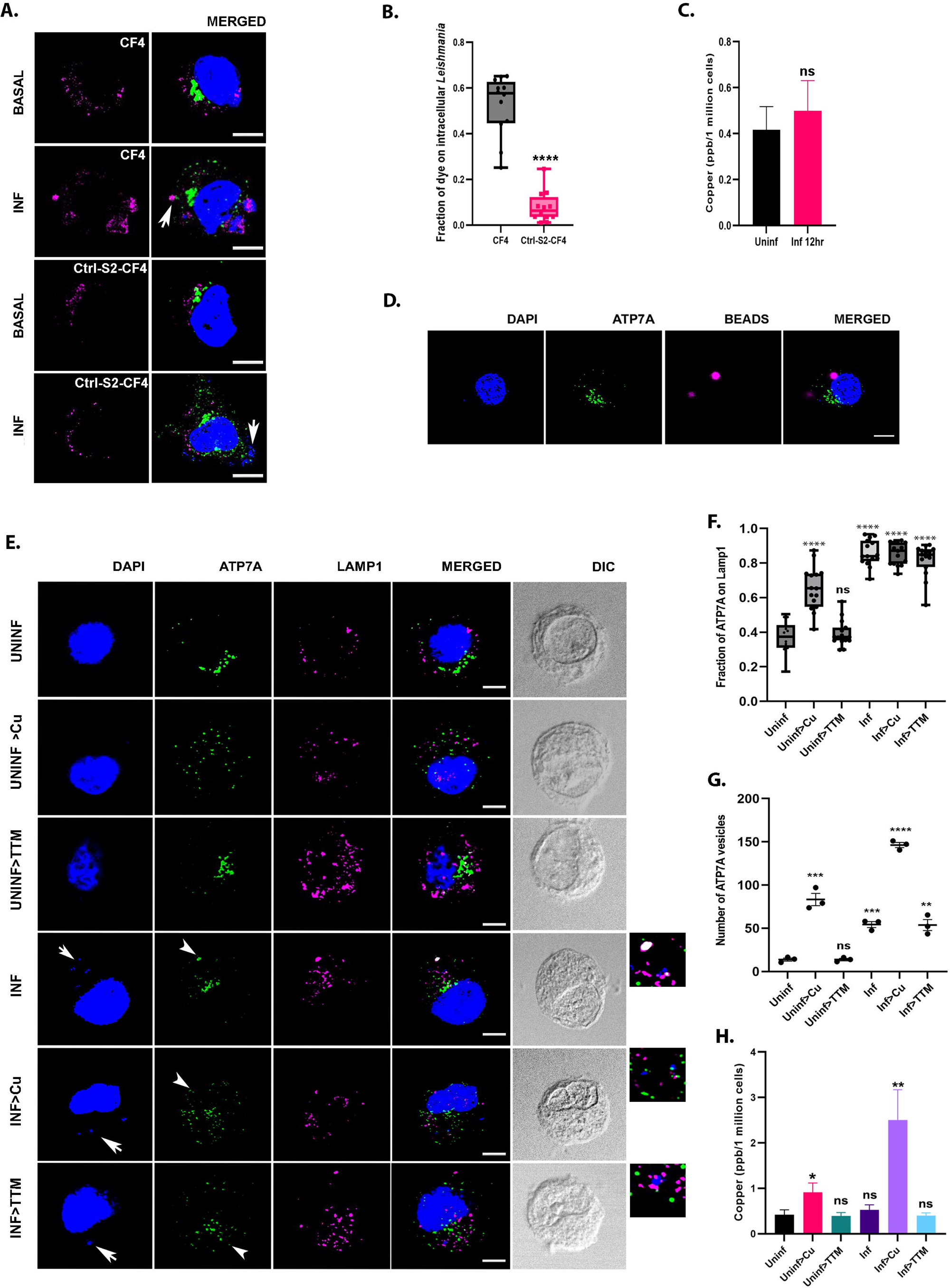
*Leishmania* infection triggers host ATP7A trafficking and copper accumulation. (A) Representative immunofluorescence image of CF4 dye or Ctrl-S2-CF4 dye (magenta), co-stained with ATP7a (green), in J774A.1 macrophages with (INF) and without (BASAL) *L.major* infection (12 hour). Both macrophage and *Leishmania* nuclei were stained with DAPI (blue). The merged images represent colocalization of the dyes with *Leishmania* nuclei. White arrows indicate intracellular parasites in infected cells (smaller nuclei). The scale bar represents 5 μm. (B) Fraction of dye colocalized with *Leishmania* nuclei from the above mentioned conditions demonstrated by a box plot with jitter points. The box represents the 25th to 75th percentiles, and the median in the middle. The whiskers show the data points within the range of 1.5× interquartile range from the first and third quartiles. Sample size (n): 12, ****p ≤ 0.0001 (Wilcoxon rank-sum test). (C) Measurement of intracellular copper level in J774A.1 macrophages with (Inf 12 hr) or without (Uninf) *L.major* infection using ICP-MS. Error bars represent mean ± SD of values calculated from three independent experiments. ns; (Student’s t test). (D) Immunofluorescence image of ATP7A (green) in J774A.1 macrophage with FluoSpheres™ Carboxylate-modified Microsphere beads (magenta) confirming absence ATP7A trafficking to those beads due to general phagoscytosis. Macrophage nuclei is stained with DAPI (blue). (E) Representative immunofluorescence image of ATP7A (green), co-stained with endo-lysosomal marker Lamp1 (magenta), in J774A.1 macrophages with and without *L.major* infection (12 hour) followed by basal, high copper (100 μM Cu) and copper chelated conditions (25 μM TTM) treatment for 2 hours. The merged images represent colocalization of ATP7A with Lamp1. Both macrophage and *Leishmania* nuclei were stained with DAPI (blue). White arrows indicate intracellular parasites in infected cells (smaller nuclei). White arrowheads indicate vesicularised ATP7A. The scale bar represents 5 μm. The magnified inset corresponds to the region of the merged image marked by arrows indicating the association of ATP7A and Lamp1 with *Leishmania*-positive endosomes. (F) Fraction of ATP7A colocalization with Lamp1 from the above mentioned conditions demonstrated by a box plot with jitter points. The box represents the 25th to 75th percentiles, and the median in the middle. The whiskers show the data points within the range of 1.5× interquartile range from the first and third quartiles. Sample size (n): 15. ****p ≤ 0.0001, ns; (Wilcoxon rank-sum test). (G) Number of ATP7A vesicles from the above mentioned conditions are plotted. (H) Comparison of copper levels (in ppb/million cells) in *Leishmania* infective uninfected cells under different treatments. Error bars represent mean ± SD of values calculated from three independent experiments. **p ≤ 0.01, ***p ≤ 0.001, ****p≤ 0.0001, ns; (Student’s t test).

*Leishmania* is phagocytosed by macrophages, subsequent to which it traverses through the endo-lysosomal compartments. To test whether macrophage ATP7A trafficking was a response to the general phagocytosis event or was specific to *Leishmania* infection, we used fluorescence beads that underwent phagocytosis. ATP7A did not traffic in response to phagocytosis of fluorescence beads (Figure 1D). ATP7A trafficking was *Leishmania* infection-specific which was further confirmed by copper treatment post-infection (Figures 1E and 1F). Macrophage J774A.1 were infected with *L. major* for 12 hours followed by 100 µM copper for 2 hours. We observed further dispersal of ATP7A in vesicles as compared to only infection (Figure 1G). Interestingly, *Leishmania* compartment-associated ATP7A localisation did not alter appreciably as there was no significant change in the colocalisation of ATP7A and Lamp1 upon copper treatment (Figure 1F). Hence, we infer that in infected cells, it was the residual Golgi fraction of ATP7A that responded to copper treatment and trafficked out. Similarly, 25 µM TTM (intracellular copper chelator) treatment for 2 hours post-infection did not cause amastigote-associated ATP7A to return to Golgi (Figures 1F and 1G). Copper and TTM treatments without infection resulted in ATP7A trafficking from and to Golgi, respectively (Figures S1A and S1B). The established role of ATP7A to vesicularize from the Golgi upon excess intracellular copper and recycle back to Golgi upon copper removal was different from infection-induced phenotype. Intracellular copper levels were estimated for these conditions by ICP-MS. Interestingly, copper level shot-up by 2.5 folds in case of post-infection copper treatment as compared to uninfected copper treatment group (Figure 1H). This observation suggest that copper uptake is augmented by the host macrophages as a mechanism to facilitate parasite killing.

### *Leishmania major* infection triggers degradation of macrophage ATP7A

The host macrophage channelizes copper in the compartments harboring *Leishmania* amastigotes as a host response towards infection. We hypothesized that a reciprocal response should be surmounted by the pathogen to circumvent host-induced copper stress to establish a successful infection. Previously, it has been shown that ATP7A protein expression increases on treating macrophages with pro-inflammatory agents known to be induced upon infection^29^ At the outset, we decided to investigate the possibilities of macrophage ATP7A modulation in infection. J774A.1 macrophages were infected with *L*. *major* and at different time points starting from 3 hours to 24 hours, the protein levels of ATP7A were determined by immunoblotting. We observed remarkable reduction of ATP7A protein during early-mid time points hours of infection (3-15 hour) that reinstated back to ∼70% of the normal level at late infection time-points (24 hour) (Figure 2A). We checked the corresponding transcript levels which remarkably were not downregulated suggesting inhibition of transcriptional machinery or RNA degradation were not contributing to low ATP7A protein levels (Figure 2B). Intracellular pathogen load heightened around 12-15 hour stage followed by reduction as evident from kDNA-based PCR method (Figure S2A). Similarly, intracellular copper level elevated around late stage of infection (15-18 hours) which corroborated with ATP7A recovery and reduced pathogen load at late stage (Figure S2B).

**Figure 2.**
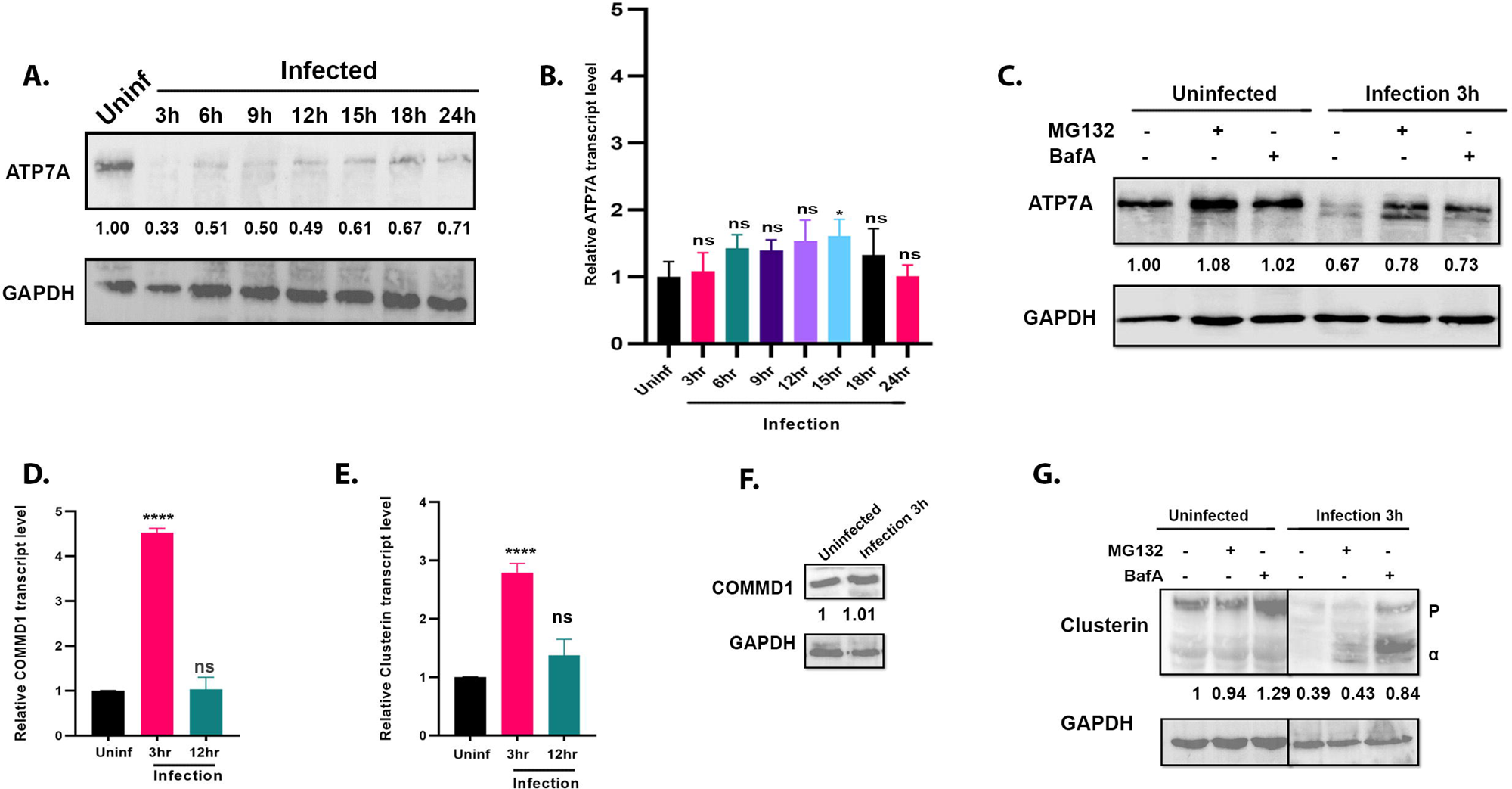
*L. major* infection causes degradation of host ATP7A by modulating regulatory proteins upstream of it. (A) Immunoblot of ATP7A at indicated time points after infecting J774A.1 macrophages with *L.major* promastigotes. The fold changes of ATP7A abundance normalized against housekeeping control GAPDH have been mentioned. (B) qRT–PCR shows ATP7A transcript level at indicated time points after infection, normalized against uninfected control and housekeeping control GAPDH mRNA levels. Error bars represent mean ± SD of values calculated from three independent experiments. *p ≤ 0.05, ns; (Student’s t test). (C) Immunoblot of ATP7A from infected and uninfected J774A.1 macrophages for 3 hours with or without co-treatment with MG132 or Bafilomycin A1. The fold changes of ATP7A abundance normalized against housekeeping control GAPDH have been mentioned. (D) qRT–PCR shows COMMD1 transcript level at 3 hour and 12 hour time points after infection, normalized against uninfected control and housekeeping control GAPDH mRNA levels. Error bars represent mean ± SD of values calculated from three independent experiments. ****p ≤ 0.0001, ns; (Student’s t test). (E) qRT–PCR shows Clusterin transcript level at 3 hour and 12 hour time points after infection, normalized against uninfected control and housekeeping control GAPDH mRNA levels. Error bars represent mean ± SD of values calculated from three independent experiments. ****p ≤ 0.0001, ns; (Student’s t test). (F) Immunoblot of COMMD1 after 3 hour infection. The fold changes of Clusterin abundance normalized against housekeeping control GAPDH have been mentioned. (G) Immunoblot of Clusterin from infected and uninfected cells with or without pretreatment with MG132 or BafilomycinA1. P denoted Clusterin precursor and α denoted Clusterin alpha chain. The fold changes of Clusterin precursor abundance normalized against housekeeping control GAPDH have been mentioned.

To determine the degradation pathway(s) of macrophage ATP7A that is triggered by *Leishmania* infection, ATP7A levels were checked by immunoblot following 3 hours infection with or without the addition of proteasome inhibitor MG132 or lysosomal inhibitor Bafilomycin A1. When compared to the uninfected control, the level of the ATP7A protein decreased after infection. Treatment with MG132 and bafilomycin A1 prevented ATP7A from degrading due to infection (Figure 2C). Therefore, when lysosomal or proteasomal inhibition was inhibited, the degradation phenotype was reversed.

It has been reported that copper stress or intracellular pathogens may induce oxidative stress^37, 38^. We tested if ATP7A degradation was a result of oxidative stress that might arise due to *Leishmania* infection or copper accumulation in amastigote harboring compartments. Macrophages were exposed to 200 µM or 500 µM of H_2_O_2_ for 2 hours to simulate oxidative stress. Similarly, 100 µM of copper for 2 hours and 10 µM of copper for 36 hours were used to simulate high and prolonged copper levels, or 50 µM of TTM (an intracellular copper chelator) for 2 hours and 10 µM of TTM for 36 hours to simulate low copper levels. Immunoblot analysis revealed no noticeable ATP7A degradation that rules out oxidative stress, copper toxicity, or copper deprivation as potential causes for the observed phenotype (Figure S2C). These results suggest that the degradation phenotype of ATP7A protein is unique to infection and may be mediated via both lysosomal and proteasomal degradation pathways. Upon stimulation by LPS and IFN-γ, ATP7A expression has been previously demonstrated to increase^29^. Interestingly, for *Leishmania* infection, we observe a novel ATP7A degradation phenotype. We speculate that this modulation is a pathogen defense mechanism to reduce Cu toxicity exerted by ATP7A, as ATP7A localizes on and pumps copper into the pathogen-harboring phagolysosomal compartments. However, upon infection, there are two fractions of ATP7A. The ATP7A pool that resides on the Golgi and the vesicularized pool that localizes on the phagolysosomal compartments. It is possible that either or both of the fractions are getting degraded upon infection.

### *L. major* infection acts upstream of ATP7A in the copper homeostasis pathway

In an effort to pinpoint the source of the protein degradation, we explored the upstream regulators of ATP7A. We focused on COMMD1 and Clusterin to identify the cause of proteasomal and/or lysosomal degradation of ATP7A. COMMD1 and Clusterin help ATP7A fold and stabilize under normal physiological by acting as chaperones for it^39, 40^. On the other hand, they facilitate degradation of mutant or improperly folded ATP7A. Overexpression of either of them has been shown to accelerate ATP7A degradation^40, 41^. Independent of one another, COMMD1 and Clusterin bind with ATP7A to produce a complex that, can be seen in a co-precipitation immunoblot and promotes degradation via various routes. COMMD1 largely mediates degradation via the proteasomal pathway, whereas Clusterin facilitates degradation via the lysosomal pathway^41^.

The transcript levels of both COMMD1 and Clusterin were determined after infecting J774A.1 macrophages with *L. major* promastigotes. By using the quantitative PCR, the transcript levels of COMMD1 and Clusterin were determined at 3 hours and 12 hours post-infection. Interestingly, after 3 hours of infection compared to the uninfected macrophage, the transcript levels of both of them were significantly higher (Figures 2D and 2E). This could enhance ATP7A degradation as a pathogen-mediated regulation through both the proteasomal and lysosomal routes^41^. Alternately, it could be the host upregulating the transcript levels in an effort to stabilize ATP7A^39^.

We examined whether the changes at the transcript are reflected in the protein levels of COMMD1 and Clusterin upon infection to further define the role of these regulators in the degradation of ATP7A. In case of COMMD1, no difference in protein levels was observed between the infected (3 hours) and control groups (Figure 2F). For Clusterin, however, there was noticeable lowering in protein levels upon infection which could be reverted by Bafilomycin A1 treatment (Figure 2G). Since Clusterin acts a chaperone for ATP7A and is undergoing degradation, potentially through lysosomal pathway, we argue that its degradation is a pathogen-mediated regulation. Absence or low amount of Clusterin leaves ATP7A vulnerable to destabilizing as well as degradation, which would be advantageous to the pathogen survival. Evidences suggest that Clusterin might be working as a pro-host protein during *Leishmania* infection. To summarize, the degradation of ATP7A is predominantly carried out via the COMMD1 mediated proteasomal route. Host-specific antibodies were checked to ensure there is no cross-reactivity with *Leishmania* (Figure S3).

### *L. major* infection induces deglycosylation of ATP7A

In addition to the degradation, infected macrophages displayed two bands of ATP7A protein in the immunoblot (Figure 2C). It has been reported that ATP7A undergoes glycosylation that stabilizes the protein. The mutant G1019D exhibits incomplete or defective glycosylation that results in a similar band profile of ATP7A in immunoblots^42^. We confirmed that the observed double bands comprise of a ATP7A glycosylated band of higher molecular weight and a non-glycosylated ATP7A band of slightly lower molecular weight as confirmed by tunicamycin treatment experiment (Figure 3A). Treatment with H_2_O_2_, Cu, or TTM as described in the previous results section, did not show similar phenotype (Figure S2C), again ruling out oxidative stress, copper toxicity, or copper shortage as the explanations for the loss or inhibition of glycosylation of ATP7A.

**Figure 3.**
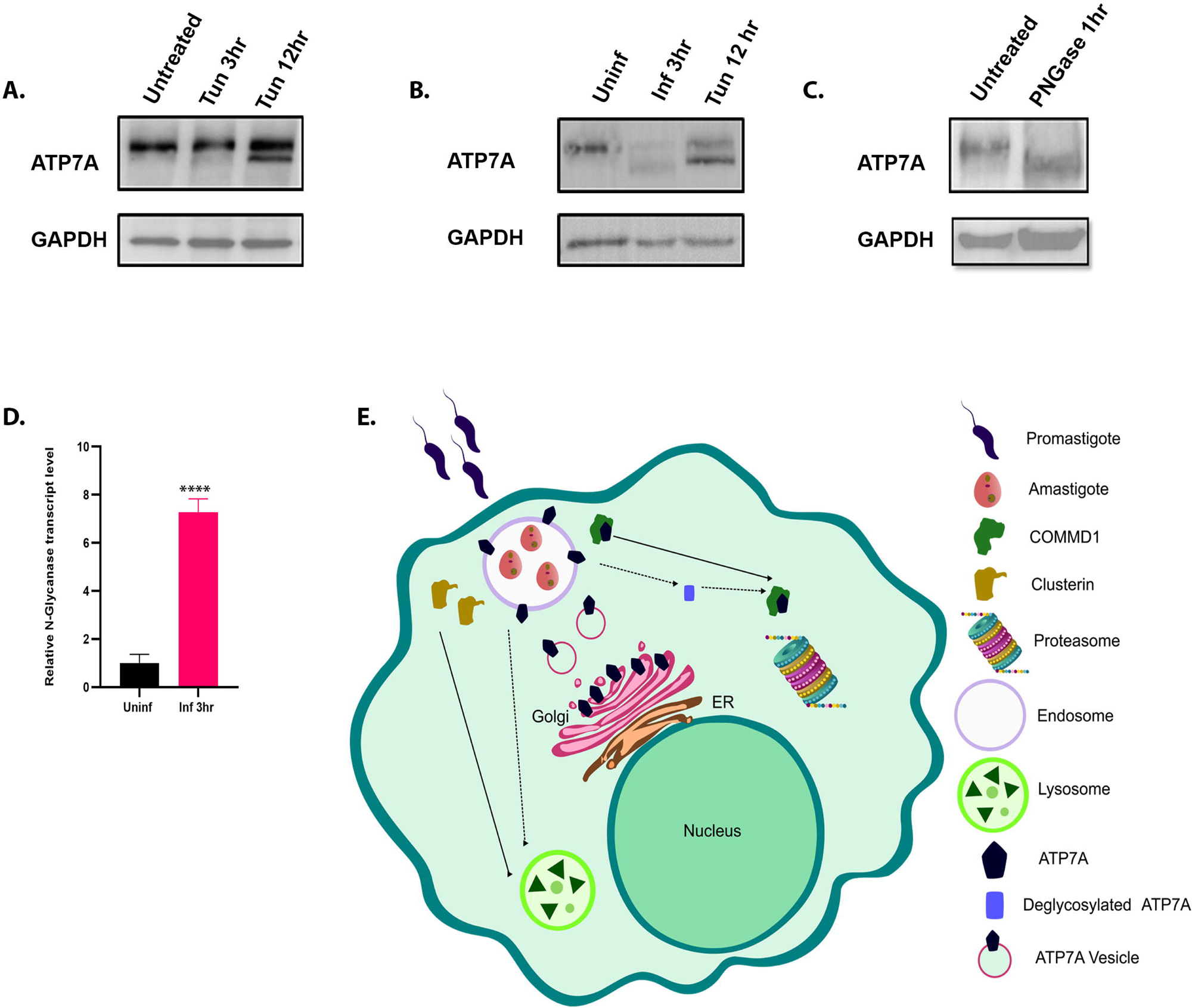
*L. major* infection triggers deglycosylation of host ATP7A. (A) Immunoblot of ATP7A at 3 hour and 12 hour time points after 1 μg/ml tunicamycin treatment of J774A.1 macrophages. GAPDH is used as a housekeeping control (B) Immunoblot of ATP7A after infecting J774A.1 macrophages with *L. major* promastigotes for 3 hours compared to 12 hour of 1μg/ml tunicamycin treatment. GAPDH is used as loading control. (C) Immunoblot of ATP7A after treatment with PNGase for 1 hour. GAPDH is used as housekeeping protein. (D) qRT–PCR shows N-G;ycanase transcript level at 3 hour time point after infection, normalized against uninfected control and housekeeping control GAPDH mRNA levels. Error bars represent mean ± SD of values calculated from three independent experiments. ****p ≤ 0.0001; (Student’s t test). (E) Illustration depicting the proposed modulation of ATP7A via the manipulation of its upstream regulators by the *L.major* pathogen.

There are two potential explanations for the appearance of this non-glycosylated band. The newly synthesized ATP7A is either not being glycosylated or the glycan chain is being removed by a pathogen-initiated process, which results in the deglycosylation of ATP7A.

To understand mechanism of loss of glycosylation, J774A.1 macrophages were treated with tunicamycin for 3 hours and 12 hours to determine if the non-glycosylated band was caused by the inhibition of glycosylation. Tunicamycin serves as an inhibitor of N-linked glycosylation for newly synthesized proteins and inhibits the reverse processes in the initial step of the manufacture of N-linked oligosaccharides in cells^43^. From the immunoblot, a very faint non-glycosylated band for ATP7A started to develop about 3 hours after tunicamycin treatment. A non-glycosylated band resembling the intensity of the one seen in infection did not appear until 12 hours post-treatment (Figures 3A and 3B). But remarkably, in infection, the non-glycosylated band can start to appear as early as 3 hours. So, it is evident that newly synthesized protein whose glycosylation has been inhibited requires a longer time period for the accumulation of non-glycosylated protein. Therefore, to explain the fast appearance of this non-glycosylated band, we hypothesized that infection is actually causing deglycosylation, or the removal of N-linked glycan chain from mature ATP7A. Infected J774A.1 macrophage cell lysates were treated with PNGaseF for 1 hour. PNGaseF is an amidase that removes nearly all N-linked oligosaccharides from glycoproteins by cleaving oligosaccharides between their innermost GlcNAc and asparagine residues. The ATP7A lower band appeared within an hour of treatment (Figure 3C). This demonstrates that deglycosylation operates more quickly than glycosylation inhibition, which may be the cause of the emergence of the non-glycosylated ATP7A upon infection.

An enzyme called N-glycanase de-N-glycosylates glycoproteins by removing N-linked or asparagine-linked glycans^44^. J774A.1 macrophages were infected with *L. major* promastigotes for 3 hours whereas uninfected macrophages served as the control. qPCR was used to determine the host N-glycanase mRNA levels in control and infected macrophages. N-glycanase transcript levels were markedly increased by 7.3 folds after infection (Figure 3D). During *Leishmania* infection, ATP7A is deglycosylated, which is possibly a direct effect of an increase in host N-glycanase. Several reports suggest interfering with the glycosylation status of proteins makes them less stable and prone to degradation^45, 46^. This could essentially make the process of ATP7A degradation easier for the pathogen.

### Knock down of ATP7A and primary copper importer CTR1 increases *Leishmania* infectivity

Upon infection, host ATP7A is manipulated by the pathogen at multiple regulatory levels (Figure 3E). The crucial role of host ATP7A in limiting parasitic load can be attributed to its ability to channel copper to *Leishmania* compartments. In mammalian copper uptake and utilization pathway, ATP7A receives copper from the high-affinity copper uptake protein 1, CTR1, facilitated by the antioxidant 1 copper chaperone, ATOX1 (Figure 4A). Degradation of ATP7A is one way of ensuring reduced copper delivery to amastigote-positive compartments. To further ascertain the importance of host ATP7A in resisting *Leishmania* infection, ATP7A was depleted using siRNA-mediated knock-down and subsequently protein levels were confirmed by immunoblotting. Macrophage treated with scrambled siRNA was used as an experimental control for infection studies. At 12 hours post-infection, amastigote burden estimation was performed by amastigote nuclei count and kDNA-based PCR method. Corroborating with our hypothesis, DAPI-stained nuclei counting revealed amastigote per macrophage was about 1.3 folds higher in ATP7A knock-down macrophages compared to the control set (Figure 4B). The Ct value of kDNA was significantly lower for ATP7A depleted infected macrophages than the control indicating higher parasite load in macrophages with downregulated ATP7A, hence conforming with our previous inference (Figure 4C). Both findings indicate that host ATP7A knockdown benefits *Leishmania*, resulting in higher infectivity.

**Figure 4.**
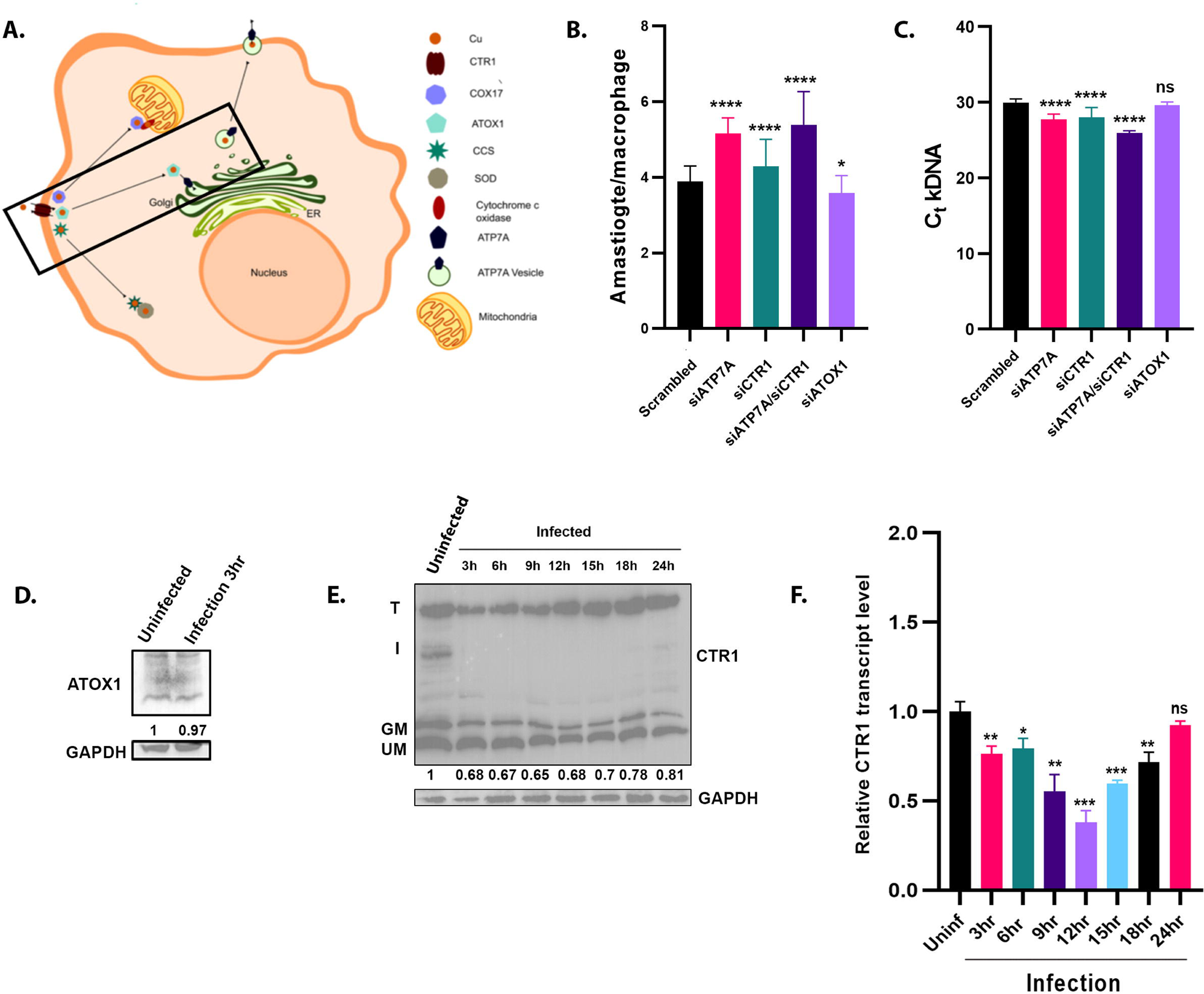
Knocking down of host ATP7A and CTR1 enhances the infectivity of the *Leishmania* pathogen. (A) Illustration depicting the mammalian copper uptake and utilization pathway. Copper secretory pathway is highlighted. (B) J774A.1 macrophages, after treatment with scrambled, ATP7A, CTR1 and ATOX1 siRNA, were infected with *L. major*. After 12 hours, amastigote counts inside the macrophage are plotted. At least 100 macrophages were counted from triplicate experiments. Error bars represent mean ± SD of values from three independent experiments. *p ≤ 0.05, ****p ≤ 0.0001; (Student’s t test). (C) J774A.1 macrophages, after treatment with scrambled, ATP7A, CTR1 and ATOX1 siRNA, were infected with *L. major*. After 12 hours, C_t_ values of *L. major* kDNA are plotted. Error bars represent mean ± SD of values from three independent experiments. ****p ≤ 0.0001, ns; (Student’s t test). (D) Immunoblot of ATOX1 at 3 hour time point after infecting J774A.1 macrophages with *L. major* promastigotes. The fold change of ATOX1 abundance normalized against housekeeping control GAPDH has been mentioned. (E) Immunoblot of CTR1 at indicated time points after infecting J774A.1 macrophages with *L. major* promastigotes. T denotes CTR1 trimer, I denotes Intermediate band, GM denotes glycosylated monomer, UM denotes unglycosylated monomer. The fold changes of CTR1 glycosylated monomer abundance normalized against housekeeping control GAPDH have been mentioned. (F) qRT–PCR shows CTR1 transcript level at indicated time points after infection, normalized against uninfected control and housekeeping control GAPDH mRNA levels. Error bars represent mean ± SD of values calculated from three independent experiments. *p ≤ 0.05, **p ≤ 0.01, ***p ≤ 0.001, ns; (Student’s t test).

Copper Transporter-1 (CTR1) or SLC31A1 is the primary copper importer in the host cell, whereas ATOX1 chaperones the copper from CTR1 to ATP7A (Figure 4A). Hence, we probed the entire copper acquisition and delivery pathway that is pre-requisite to ATP7A functioning. siRNA-based knock-down of both CTR1 and ATOX1 in macrophage followed by infection studies were performed to ascertain the role of these key copper regulators in resisting *Leishmania* infectivity. siRNA knock-downs were confirmed by immunoblotting (Figure S4). CTR1-depleted cells had low kDNA Ct value and high amastigote per macrophage count compared to control (scrambled siRNA-treated macrophage) (Figures 4B and 4C). Similar to ATP7A, CTR1 seemed to have a resistant role to *Leishmania* infection. Interestingly, no such alteration in infection levels was found in ATOX1 knock-down infected macrophages.

CTR1 and ATP7A are key candidates in maintaining cellular copper homeostasis. CTR1 knock-down resulted in altered copper uptake which compromised its delivery and availability to ATP7A. This resulted in an overall decrease in copper-induced toxicity on intracellular amastigotes. When both CTR1 and ATP7A were depleted together in the host macrophage, infection increased further by 1.39 folds, as evident from amastigote nuclei count and kDNA Ct value (Figures 4B and 4C). The ability of CTR1 to control the copper intake and accumulation, in addition to the directionality of copper deployment provided by ATP7A towards amastigote-harboring compartments, could be key factors in *Leishmania* infection and survivability.

To study if *Leishmania* manipulates host copper uptake and distribution to evade host response, we probed the protein levels of ATOX1 and CTR1 upon infection. We did not observe an alteration in ATOX1 protein amount (Figures 4D). In immunoblots, CTR1 is found in three bands possibly representing the three distinct oligomeric forms of the protein, (a) mature functional trimer (∼100kDa), (b) a glycosylated monomer (∼34kDa) and (b) an intermediate form (∼65kDa) possibly representing a dimeric form that arises from a loose-trimeric state. An unglycosylated monomeric for of CTR1 (25kDa) is also observed. Interestingly for CTR1, we observed an overall lowering in the protein amount upon initial hours of *Leishmania* infection (3-12 hours). Similar to ATP7A, CTR1 protein amount restores back at late stages of infection (18-24 hour) (Figure 4E). Using RT-PCR, we determined that this reduced amount in protein levels were a direct effect of decreased CTR1 transcript upon infection (Figure 4F).

Remarkably, the mechanism of lowering CTR1 protein level was distinct from the copper regulatory protein ATP7A. Apart from the transcriptional downregulation, we observe the disappearance of the intermediate CTR1 band (65kDa) in the immunoblot of the infected macrophages. It would be interesting to find the cause and ramifications of such changes in copper importer CTR1.

### CTR1 endocytose upon *Leishmania* infection

CTR1 is the primary copper importer in the cell. Under basal copper levels, CTR1 localizes at the plasma membrane where it carries out its copper import function. In excess copper inside the cell, as a self-regulatory mechanism to limit copper uptake, it endocytoses from the plasma membrane via Clathrin-mediated endocytosis. When copper levels return back to normal, CTR1 recycles back to the plasma membrane^4^.

Interestingly, macrophage CTR1 endocytosed upon *Leishmania* infection (Figures 5A). We ruled-out the possibility that a general trigger of phagocytosis does not induce CTR1 endocytosis by using fluorescent beads (Figure S5). The phenotype of CTR1 endocytosis upon *Leishmania* infection is similar to when macrophages are treated with copper. Similar to ATP7A trafficking studies, we subjected *Leishmania*-infected macrophages to 2 hours of 100 µM copper and 25 µM TTM treatment separately. For the copper treated one, further CTR1 endocytosis was observed. For the TTM treated cells, we observe a slight reduction in CTR1 endocytosis (Figure 5B). However, there was a significant amount of endocytosed CTR1 in TTM treated infected macrophages, suggesting that CTR1 were not responding to copper levels; rather the observed phenotype is infection-specific. Further, immunoblot of CTR1 from macrophages under copper treatment or copper chelation was not devoid of intermediate band unlike the case during infection (Figure 5C). Nonetheless, CTR1 endocytosis (resulting in its removal from the plasma membrane) during infection will result in reduced copper uptake. Subsequently along the copper utilization pathway, ATP7A will have less available copper to mediate its targeting to pathogen compartments.

**Figure 5.**
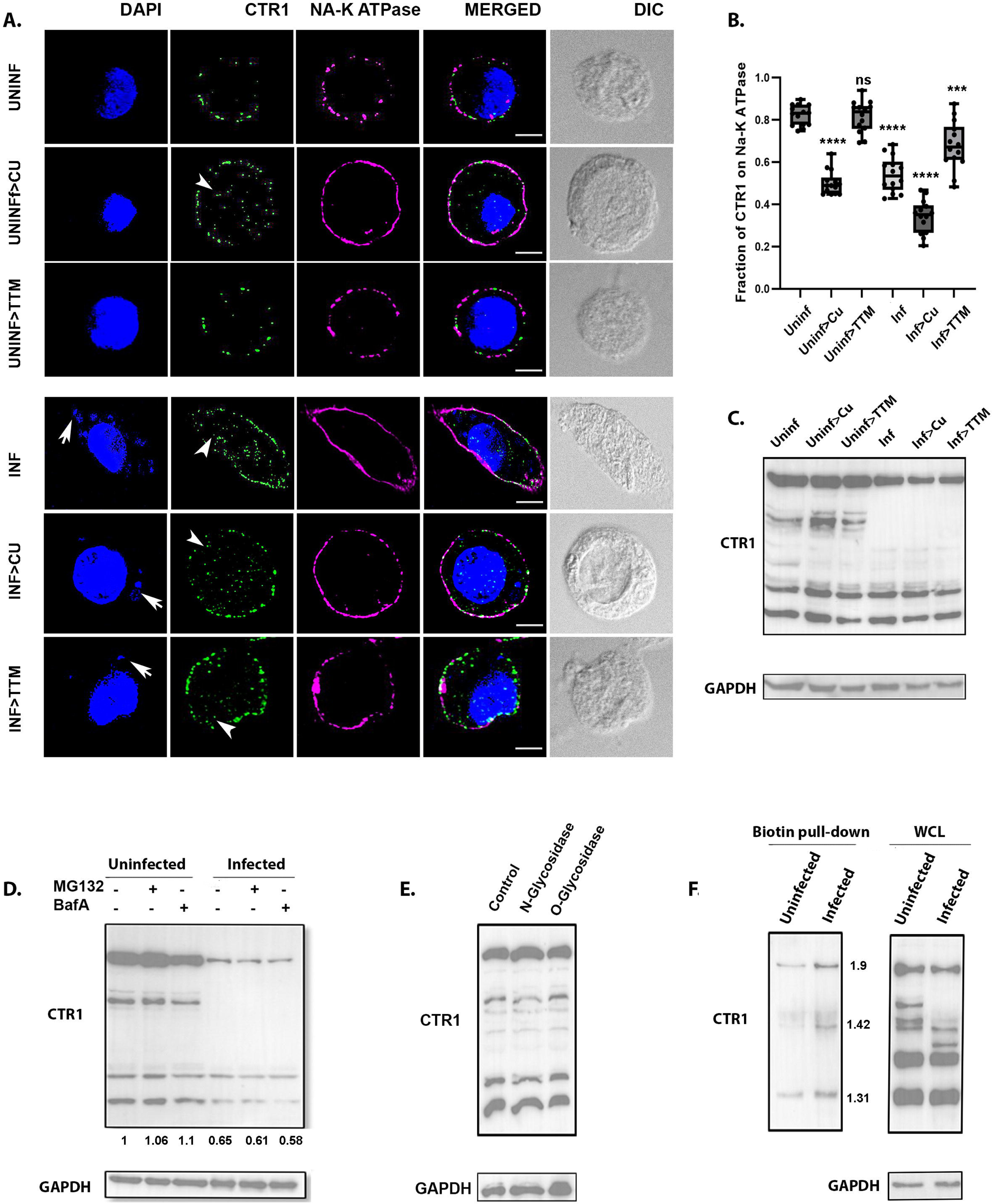
*Leishmania* manipulates host CTR1 in multiple ways to reduce copper import. (A) Representative immunofluorescence image of CTR1 (green), co-stained with plasma membrane marker Na-K ATPase (magenta), in J774A.1 macrophages with and without *L. major* infection (12 hour) followed by basal, high copper (100μM Cu) and copper chelated conditions (25μM TTM) treatment for 2 hours. The merged images represent colocalization of CTR1 with Na-K ATPase. Both macrophage and *Leishmania* nuclei were stained with DAPI (blue). White arrows indicate intracellular parasites in infected cells (smaller nuclei). White arrowheads indicate endocystosed CTR1. The scale bar represents 5 μm. (B) Fraction of CTR1 colocalization with Na-K ATPase from the above mentioned conditions demonstrated by a box plot with jitter points. The box represents the 25th to 75th percentiles, and the median in the middle. The whiskers show the data points within the range of 1.5× interquartile range from the first and third quartiles. Sample size (n): 14, ***p ≤ 0.001,****p≤ 0.0001, ns; (Wilcoxon rank-sum test). (C) Immunoblot of CTR1 of J774A.1 macrophages with or without infection (12 hour) followed by indicated copper conditions. GAPDH is used as housekeeping control. (D) Immunoblot of CTR1 from infected and uninfected J774A.1 macrophages for 3 hours with or without co-treatment with MG132 or Bafilomycin A1. The fold changes of CTR1 glycosylated monomer abundance normalized against housekeeping control GAPDH have been mentioned. (E) Immunoblot of CTR1 of J774A.1 macrophages after N-Glycosidase (PNGase 1hour) or O-glycosidase (O-Glycosidase & Neuraminidase Bundle 1hour) treatment. GAPDH is used as loading control. (F) Immunoblot of CTR1 from elute of biotin-streptavidin based pulldown of modified cysteine containing proteins from J774A.1 macrophages; and whole cell lystaes, with or without infection for 3 hours. Fold changes of CTR1 in infected samples of the elute of the biotin pulldown normalized against uninfected control have been mentioned. GAPDH is used as housekeeping control for whole cell lysate.

### *Leishmania* infection alters the redox status of CTR1 cysteines

We tried to understand the mechanism by which *Leishmania* manipulates host CTR1 via the loss of its intermediate band. Blocking the proteasomal and lysosomal degradation pathway by MG132 and Bafilomycin A1 respectively during infection were not effective to bring up the CTR1 levels and make the intermediate band reappear (Figure 5D). This further reconfirms that transcription downregulation contributed to lowering of CTR1 levels (shown in Figure 4F).

CTR1 undergoes N-linked and O-linked glycosylation^47, 48^. Since glycosylation status of host ATP7A seems to have been compromised during *Leishmania* infection, it is possible that intermediate CTR1 band might be susceptible to such actions. To check this, infected macrophage lysates were treated with deglycosylating agents. PNGase was used to remove N-glycans and O-Glycosidase and Neuraminidase bundle to remove O-glycans from CTR1. These lysates were then observed for CTR1 on immunoblot where no apparent difference or distribution in the band pattern was seen between them and uninfected macrophage CTR1 (Figure 5E). The intermediate band persisted upon inducing deglycosylation.

It has been reported that pathogenic invasion induces oxidative stress in macrophages^38^. Cysteine oxidation of CTR1 has been reported upon VEGF stimulation but not copper treatment^49^. To test whether Cys residue form Cys-sulphenic acid during infection that is an intermediate while forming the -S-S- (oxidised) form, we labelled Cys of macrophage by DCP-Bio1 (Merck). DCP-Bio1 detects Cysteinsulphenic acid and because of biotin associated with it, it can pull-down all modified cysteines by streptavidin beads. From immunoblot, we compared CTR1 from lysates of infected vs uninfected cells and observed increased pulldown of cysteine oxidised CTR1 in infected macrophages (Figure 5F). Cysteines of CTR1 have been shown to assist in dimer formation^47, 50^ and alteration in their redox state upon infection manifest into disappearance of the band. These changes apparently seem to work in favour of the parasite with the ultimate aim to reduce the amount of copper available in the cell.

### Copper bioavailability in host is critical to fight off *Leishmania* infection *in vivo*

Macrophage deploys the intracellular copper store as a mechanism to neutralize *Leishmania* infection. Reciprocally, to evade host response, *Leishmania* infection alters two key candidates of the copper secretory pathway, CTR1 and ATP7A. This manipulation is an effort to reduce the amount of copper faced by intracellular pathogen.

We checked the effect of copper availability upon *Leishmania* infection using BALB/c mice as hosts. BALB/c were categorised in three groups; first group was control (fed with regular diet and water), second group was fed with copper (20mg/L in drinking water) and third group was gavaged daily with the copper chelator, TTM (5mg/kg). After 4 weeks, serum copper was measured from each group. Copper treated group were having significantly high serum copper as compared to TTM treated and control groups (Figure S6A). To assess the efficacy of TTM as a copper chelator, we checked bioavailable serum copper level by performing serum ceruloplasmin assay^51^. Serum samples from each group were used to oxidise equal amount of O-dianisidine for a fixed period of time followed by spectrophotometric reading at 540 nm. Copper containing Holo-ceruloplasmin levels, calculated from its enzymatic activity, was used as readout of bioavailable copper^52^. Copper treated serum samples had higher ceruloplasmin activity than controls owing to high bioavailable copper. The control group had high bioavailable copper than TTM treated group indicating that TTM was indeed chelating systemic copper (Figure S6B). Copper deficiency has been reported to cause anaemia^53^ which makes it important to use low but optimal amount of TTM. Haemoglobin estimation from each group revealed no such significant changes in its level suggesting that treatment condition is optimal and could be continued and infection could be introduced in these groups (Figure S6C). These groups were further divided into two categories, uninfected and infected.

Infection was performed by subcutaneous injection to the left hind footpad of BALB/c mice with 10^6^ *Leishmania major* promastigotes. The onset and progression of lesion development in these groups of mice was monitored up to week 15 post-infection^8^. Serum total copper (including bioavailable copper) was estimated after 4 weeks of infection from each of the six groups (Uninfected, Infected, Copper, Copper Infected, TTM Infected). At four weeks post-infection, although there was no onset of lesion in the infected groups, serum bioavailable copper increased for all cases of infection (Figures 6A and 6B). It was an early indication that in response to the footpad resident *L. major*, the systemic copper level has increased to combat infection. TTM Infected group were the first one to develop visible lesion at week 6 post-infection. In the following week, there was onset of lesion in only ‘Infected group’. Interestingly, Copper Infected group showed no noticeable lesion by then, and it took week 12 to develop lesion. The trend of lesion development in all three groups was observed till week 15 that exhibited heightened lesion development in TTM Infected group (Figures 6C and D). Further, Copper Infected group had low parasite load as evident from kDNA C_t_ value (Figure 6E). Serum copper and its bioavailable form was again measured at the end of the experiment at week 15. We found similar increase in bioavailable copper in all cases of infection (Figures 6F and 6G). Naturally in Copper Infected group, overall serum copper increased that was further reflected in increased bioavailable copper which dampened the infection. It is interesting to note that even though copper levels remained high in the systemic circulation of this group, infection triggered a further increase in copper indicating possible release of copper from storage organs (Figures 6F and 6G).

**Figure 6.**
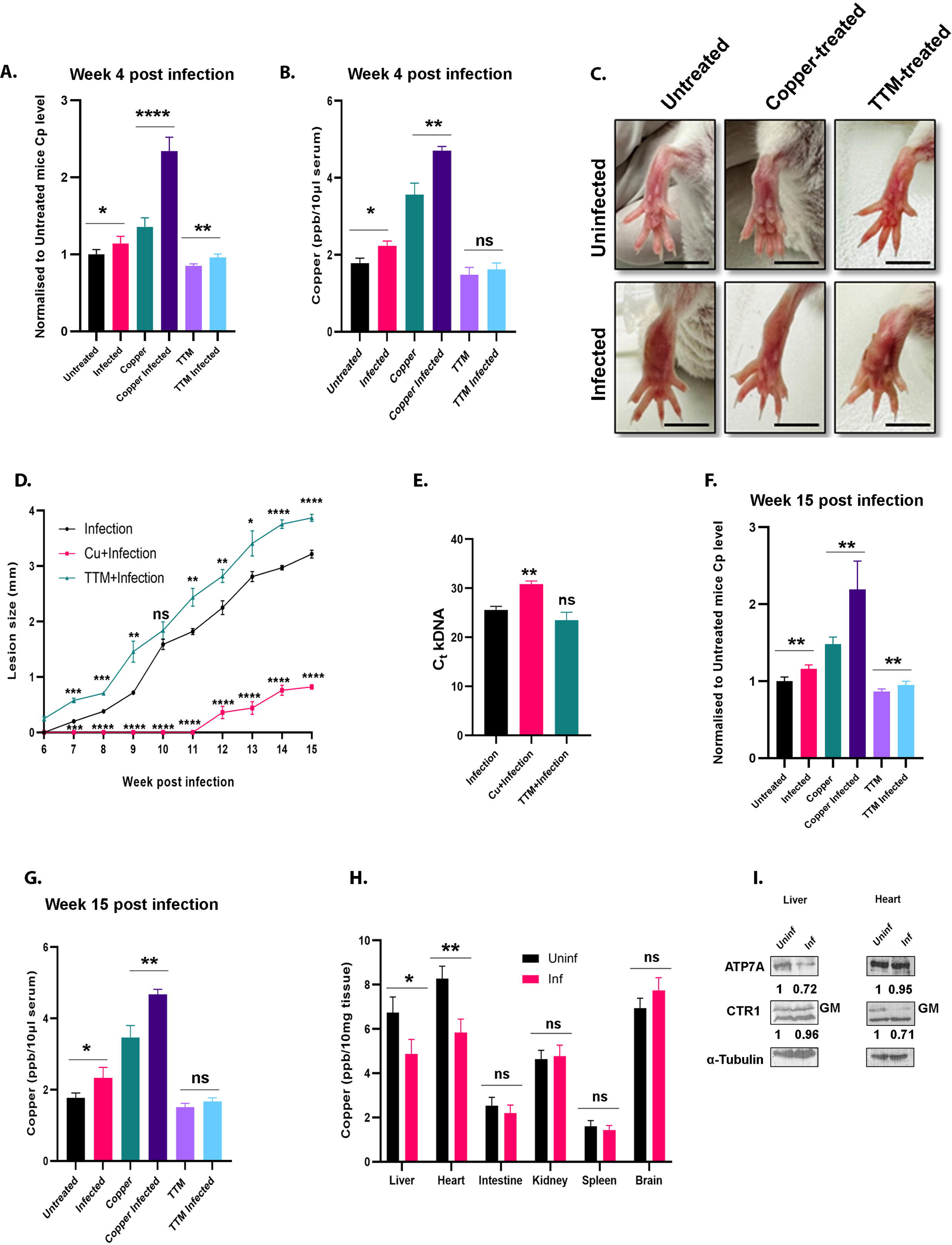
Increased bioavailable copper dampens *Leishmania* infection in vivo. (A) Measurement of Ceruloplasmin (Cp) level in serum samples of mice from indicated treatment conditions and normalised to that of the untreated mice and (B) whole serum copper from the same samples using ICP-MS at 4 weeks post-infection. Error bars represent mean ± SD of values. Sample size (n): 3 for each condition, *p ≤ 0.05, **p ≤ 0.01, ****p ≤ 0.0001, ns; (Student’s t test). (C) Represenattive images of lesion development in left footpad of infected and uninfected BALb/c mice at different copper treatments at 15 weeks post-infection. The scale bar represents 0.5 cm. (D) Line graph with markers representing progression of lesion development and size in footpad of BALB/c mice infected with *L. major* at different copper treatments. Lesion development was determined by weekly measuring the swelling with a caliper. The data correspond to the mean ± SD of values obtained from three individual mice in each group. *p ≤ 0.05, **p ≤ 0.01, ***p ≤ 0.001,****p≤ 0.0001, ns; (Student’s t test). (E) *Leishmania* load in the infected footpad was determined by Ct values of *L. major* kDNA of infected mice 15 weeks post-infection at different copper conditions. Sample (n): 3 from each group, **p ≤ 0.01, ns; (Student’s t test). (F) Measurement of Ceruloplasmin (Cp) level in serum samples of mice from indicated treatment conditions and normalised to that of the untreated mice and (G) whole serum copper from the same samples using ICP-MS at 15 weeks post-infection. Error bars represent mean ± SD of values. Sample size (n): 3 for each condition, *p ≤ 0.05, **p ≤ 0.01, ns; (Student’s t test). (H) Measurement of copper levels in the liver, intestine, heart, kidney, spleen and brain from infected and uninfected groups of mice using ICP-MS at 15 weeks post-infection. Error bars represent mean ± SD of values. Sample size (n): 3 for each condition, *p ≤ 0.05, **p ≤ 0.01, ns; (Student’s t test). (I) Immunoblot of ATP7A and CTR1 from liver and heart tissue of infected and uninfected mice. The fold changes of ATP7A and CTR1 glycosylated monomer abundance normalized against housekeeping control α-tubulin have been mentioned.

### Inhibition of cardiac copper uptake elevates systemic copper to combat infection

To identify the organ(s) that might be responsible to release copper into systemic circulation to combat infection, we compared the copper status in different organs from uninfected and infected groups. Reduced copper levels were observed in heart and liver in case of Infected group (Figure 6H). As a host response, this lowering of copper is possibly leading to enhanced systemic copper that is now channelized towards fighting-off the infection. Previously, it has been reported that copper status of the heart could signal copper mobilisation from other organs like liver and intestine^54^. We speculate that upon infection, a similar signal is generated which demands increased systemic copper to combat infection.

To understand the mechanism of channelizing heart copper towards systemic circulation, we measured the abundance of proteins responsible for copper uptake (CTR1) and efflux (ATP7A) in the heart. Interestingly, we observed that heart CTR1 protein level decreased in Infected groups which could account for low cardiac copper level (Figure 6I). As reported earlier, this would result in increased systemic copper as observed in this case. Interestingly, liver copper level also reduces which adds to the systemic copper. However, we do not observe any upregulation of its ATP7A which is involved in copper mobilisation (Figure 6I). We propose that *Leishmania* infection triggers a signalling that causes low cardiac copper and reduced heart CTR1 levels, which in turn signals for copper mobilisation from liver to the systemic circulation. It is also possible that liver and heart are signalled directly to channelize copper upon infection. Similar to the cell line data, regulation of systemic and organ copper levels is perturbed upon *Leishmania* infection. The host makes a concerted effort to increase systemic copper that seems serves as the host response to combat infection.

## Discussion

Copper is indispensable for the survival of all eukaryotic life forms. However, excess copper is toxic in nature and requires tight regulation in our body. Toxic nature of copper has been utilized as an antimicrobial agent since times immemorial. Copper has been found to play a pivotal role in host-pathogen interaction^2, 28, 29^. As hosts evolve to develop resistance against pathogens, pathogens too evolve along with the host to combat host defence response. One of the most important defence cells of the innate immune system are macrophage cells, which mediate the initial response to a variety of pathogens, in an attempt to eliminate infection.

*Leishmania* parasites however, not only infect macrophages specifically, but are able to survive and thrive as intracellular amastigote form. Previously, our group has shown host copper transporting ATPase ATP7A localizes to the *Leishmania* containing phagolysosomal compartments upon infection and possibly pumps copper into these compartments to exert copper toxicity. Pathogen combats this toxicity by upregulating the expression of their own copper exporter *LmATP7*. In the present study, we identified a novel parasite-induced trafficking phenomenon of the host ATP7A which is independent of copper cues. We speculated that ATP7A might have an important role in host defence against *Leishmania* pathogenesis.

Macrophage cells also express copper transporters and copper regulating and interacting proteins, which can play an important role in host defence to exert copper toxicity on the pathogen. It has been reported that ATP7A protein expression increases on treating macrophages with pro-inflammatory agents that are elevated in microbial infection^29^. However, we observe that upon infection of macrophages with *L. major* promastigotes, there is a remarkable degradation of ATP7A during early stages of infection. It would be interesting to explore the proteome change in the host and the parasite at different time points of infection to pinpoint possible soluble factor(s) that might signal trafficking of ATP7A at early time points and its degradation subsequently.

Degradation of ATP7A during microbial infection is a unique phenomenon and has not been reported earlier. Interestingly, ATP7A degrades via both the proteasomal and lysosomal degradation pathways. COMMD1 and Clusterin are two regulators of ATP7A. Under physiological conditions, they act as chaperones and interact with ATP7A to form a complex to help with proper folding of ATP7A and stabilize it. However, when ATP7A is misfolded beyond repair, COMMD1 and Clusterin independently associate with ATP7A to target it towards proteasomal and lysosomal degradation respectively^41^. We observed that the transcript levels of COMMD1 and Clusterin were both elevated upon infection which support the idea that ATP7A is getting degraded via both the proteasomal and lysosomal pathways upon infection.

Upon exploring the possible reasons of reduced stability of macrophage ATP7A in *Leishmania* infection, we found that infection induces deglycosylation of the protein as observed by appearance of both a glycosylated and a non-glycosylated band in that experimental condition. This observation was particularly interesting as previous studies show that loss of glycosylation leads to reduced functional activity of a protein^45^ and also decreases stabilization of ATP7A on membranes^46^. This suggests that the loss or inhibition of glycosylation of ATP7A is facilitating the degradation of ATP7A by the parasite. We for the first-time report that *Leishmania* upregulates the host N-glycanase, a key enzyme that remove sugar residues from mature glycoproteins. Our observation reveals deployment of a multi-modal manipulation of the host copper homeostasis pathway by *Leishmania*.

Nevertheless, it seems that primarily the phagolysosomal fraction of ATP7A, which is directly exposed to the pathogen, is undergoing the modulations and not the Golgi residing fraction. As mentioned previously, a considerable fraction of ATP7A remains at the Golgi after infection and respond to copper cues as well. Whereas, the phagolysosomal fraction that responded to infection fails to respond to further copper cues. Figure 3E illustrates a model based on our findings describing the role of ATP7A and its regulators in *Leishmania* infection.

*Leishmania* uses multiple strategies to modulate host ATP7A to make it less potent. ATP7A receives copper from High affinity copper uptake protein 1, CTR1. In the plasma membrane, CTR1 uptakes copper which is taken up by the metallochaperone, ATOX1 and transferred to ATP7A. siRNA-mediated knockdown study revealed ATP7A and CTR1 are crucial players in restricting intracellular infection indicating both copper intake and channelization are pivotal in the process of host-response.

Like ATP7A, CTR1 too was modulated by *Leishmania*. Interestingly, the strategy employed here by the parasite is distinct from the one for ATP7A. CTR1 was transcriptionally downregulated at early stages of infection. We confirmed that unlike ATP7A, CTR1 was not subjected deglycosylation or degradation. Interesting, CTR1 from infected macrophages underwent Cys oxidation which could be one of the plausible reasons for the disappearance of the intermediate bands. Previous reports suggest involvement of Cys of CTR1 in dimer formation. It is evident that transcriptional downregulation of CTR1 is beneficial for pathogen as functional CTR1 adds to the potency of ATP7A. It will be interesting to investigate how the loss of intermediate CTR1 band effect its functionality and whether its beneficial for the host or pathogen. We hypothesize based on our findings (unpublished data) that the appearance of the CTR1 dimer in a reducing gel is a reflective of the copper import capacity of the transporter. Disappearance of the same is possibly indicative of compromised copper uptake by CTR1.

We extended our study to the well-established mouse model system where we tweaked with copper bioavailability in mouse. Copper treated and TTM treated groups having high and low bioavailable copper respectively compared to untreated group were subjected to *L. major* infection. Copper treated group had developed small lesions at a later time point of infection. TTM group were the first one to develop lesion and maintained a heightened lesion size and parasite load till the end of the infection as compared to the control and copper treated groups. Interestingly, serum copper level including its bioavailable form increased in all the groups upon infection. This indicates that introducing *Leishmania* on the mice footpad had channelised copper into the systemic circulation and finally to the infection site. Upon investigating the source of elevated copper in systemic circulation, we found that heart acts as the major copper donor by reducing its copper uptake by downregulating CTR1. It has been demonstrated previously by Kim and coworkers that in mice with heart-specific knockout of CTR1, accumulated less copper as compared to wild-type littermates. Consequently, hepatic copper is decreased by approximately 20%, and serum Cu is elevated by approximately 30%. These data corroborate with our finding that heart copper is diverted towards systemic circulation for its utilization to fight-off infection. Similarly, in the hepatic tissue we noticed a decrease in copper levels indicative of hepatic copper being utilized as well in systemic circulation. However, based on immunoblots of hepatic ATP7A is neither elevated or CTR1 downregulated in infection condition. We hypothesize that liver copper is channelized to systemic circulation possibly via other liver-specific metal transporter that calls for further investigation.

Using the model of host-parasite interaction, our study also helps to understand better the intricacies of the mammalian copper metabolism pathway at a systemic level. We for the first time show an interplay of the copper importers and exporters/ATPases at a tissue level maintaining an optimum level of systemic copper at a given physiological or pathophysiological situation.

To summarize we propose a model of host-*Leishmania* infection where both partners employ various means to attune the copper homeostasis pathway in a way that serves towards their respective advantage (Figure 7). At early time points of infection, the parasite establishes a successful infection by downregulation the two key host copper transporters. However, at a later stage of infection, the host responds by channelizing its copper stores to the site of infection to attenuate the parasite.

**Figure 7.**
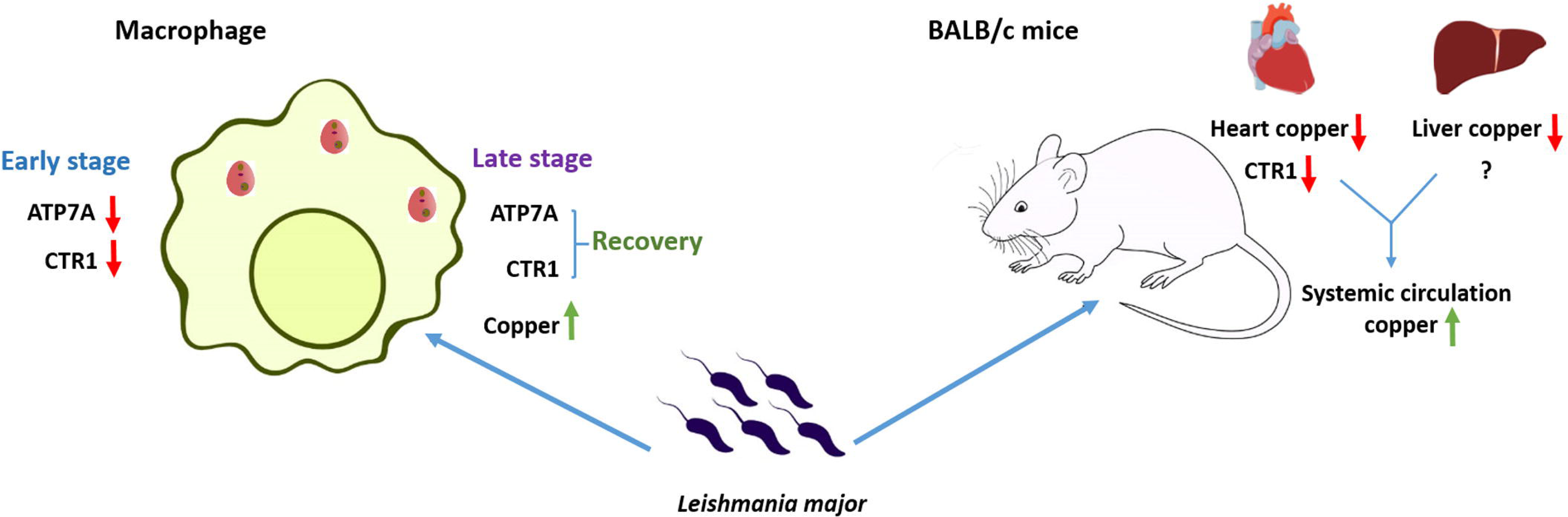
Proposed model depicting the alteration of copper homeostasis pathway at cellular and systemic level upon *Leishmania major* infection Macrophage ATP7A and CTR1 face early reduction upon *Leishmania* infection but recover back to exert copper stress on pathogen. In BALB/c mice, infection results in increased systemic copper level, contributed by heart and liver, to fight the pathogen.

## Materials and Methods

### Ethics Statement

The animal studies involving mice were approved by IISER Kolkata Animal Ethics Review Board and experiments were conducted according to the Committee for the Purpose of Control and Supervision of Experiments on Animals guidelines (CPCSEA) and Institutional Animal Ethics Committee (IAEC) approved protocol. BALB/c mice were obtained from the National Institute of Nutrition (NIN), Hyderabad, and were housed in our institutional animal facility.

### Parasite and mammalian cell culture

*L. major* promastigotes strain 5ASKH (a kind gift from Dr Subrata Adak, Indian Institute of Chemical Biology) were grown in M199 medium (Gibco #11150067) supplemented with 15% FBS (Gibco #10270106), 23.5 mM HEPES, 0.2 mM adenine, 150 μg/ml folic acid, 10 μg/ml hemin, 120 U/ml penicillin, 120 μg/ml streptomycin, and 60 μg/ml gentamicin at pH 7.2, and the temperature was maintained at 26 °C. The murine macrophage cell line, J774A.1 (obtained from the National Centre for Cell Science), was grown in DMEM (Sigma #D6429) supplemented with 2 mM l-glutamine, 1X penicillin-streptomycin, and 10% FBS at 37 °C in a humidified atmosphere containing 5% CO2.

### Macrophage infection with *L. major* and other treatments

J774A.1 macrophages were infected at a density of 1 × 10^6^ cells with late log-phase *L. major* promastigotes as described previously at a parasite-to-macrophage ratio of 30:1^8^. Infected macrophages were incubated for specific time-points following which the cells were washed with twice 1X PBS to remove promastigotes from the medium. They were either subjected to further treatments or harvested to carry out various studies. J774A.1 macrophages were treated with 10µM MG132 (Sigma) and 0.16µM Bafilomycin A1 (Sigma) for 3 hours with or without infection following which the cells were harvested for immunoblotting. Macrophages were harvested for immunoblotting after treatment with 200µM or 500µM of H_2_O_2_ (Sigma) for 2 hours, 100µM and10µM of copper (Sisco Research Laboratories) for 2 hours and 36 hours respectively, and 50µM and 10µM of TTM (an intracellular copper chelator) (Sigma) for 2 hours and for 36 hours respectively.

### Immunofluorescence and microscopy

J774A.1 macrophages were seeded at a density of 2 × 10^5^ cells on glass coverslips. After following different experimental set ups to required time points, cells were fixed using chilled acetone:methanol (1:1) for 15 min by keeping on ice. Cells were then washed with ice-cold 1X PBS and blocked with 3% BSA in PBSS for 1 hour at room temperature. Antibody solution was made in 1% BSA in PBSS. Cells were incubated in primary antibody solution (anti-ATP7A 1:300; anti-Lamp1 1:200; anti-CTR1 1:500; anti-Na-K ATPase 1:500) for 2 hours at room temperature followed by three 1X PBS wash. Cells were then reincubated for 1.5 hours at room temperature with secondary antibody solution (goat anti-rabbit Alexa Fluor 488 at 1:1000 for ATP7A and CTR1, goat anti-mouse Alexa Fluor 568 (1:1000) for Lamp1 and Na-K ATPase). Following three 1X PBS washes, coverslips were mounted on glass slides using 10µL of Fluoroshield with DAPI mountant (Sigma #F6057). All images were captured with Leica SP8 confocal platform using oil immersion 63× objective and were deconvoluted using Leica Lightning software. Antibody details are listed in Table S2.

### Latex bead phagocytosis in Macrophages

J774A.1 macrophages were incubated for 12 hours with 1 μM FluoSpheres™ Carboxylate-modified Microspheres, red fluorescent (580/605) (Molecular Probes, Life Technologies) (Kind gift of Dr. Bidisha Sinha, IISER Kolkata). Following incubation, cells where fixed and processed for immunofluorescence studies as mentioned earlier.

### Imaging of cellular labile copper with CF4 dye

J774A.1 macrophages with or without infection (12 hour) were treated either with 0.8 μM CF-4 or control CF-4 (Ctrl-S2-CF4) in DMEM without phenol red (Gibco) and incubated for 10 min at 37°C and 5% CO_2_. Following incubation, cells are washed twice with DMEM without phenol red to remove excess probe and were kept for an additional 20 min at 37°C and 5% CO_2_. Cells were fixed and processed for immunofluorescence studies as mentioned earlier.

### Immunoblotting

After respective treatments with infection, J774A.1 macrophage cells were pelleted down. Cells were resuspended in RIPA lysis buffer (10mM Tris-Cl pH 8.0, 1mM EDTA, 0.5mM EGTA, 1.0% Triton X-100, 0.1% sodium deoxycholate, 0.1% SDS, 140mM NaCl, 1X protease inhibitor cocktail, 1mM phenylmethylsulfonyl fluoride) and kept for 15 mins on ice. For immunoblotting from tissue, tissue was collected from euthanized mice following transcardial perfusion with PBS followed by flash-freezing in liquid nitrogen. 100 mg tissue for liver, heart, intestine, kidney, spleen and brain was lysed with 1 ml RIPA lysis buffer. The solution was then sonicated with a probe sonicator (4 pulses, 5s on, 20s off and 100mA). Protein sample were quantified using Bradford reagent (Sigma) and 20μg protein was loaded in each well. 4X NuPAGE loading buffer (Invitrogen #NP0007) was added to the sample to make a final concentration of 1X and ran on 8% SDS PAGE till resolved to desired extent. After that semi-dry transfer of proteins was performed to transfer the proteins from blot to nitrocellulose membrane (Milipore #IPVH00010). Following transfer, the membrane was blocked with 3% BSA in TBST for 2 hours at room temperature with mild shaking. Membranes containing proteins were incubated with primary antibody (anti-ATP7A 1:1000; anti-COMMD1 1:2000; anti-Clusterin 1:1000; anti-CTR1 1:5000; anti-GAPDH 1:3000: anti-α-Tubulin 1:10000) overnight at 4°C with mild shaking and then washed with 1X TBST. HRP conjugated respective secondary incubation was performed for 1.5 hours at room temperature, further washed with TBST and TBS. Chemoluminicent signal was developed by Clarity Max Western ECL Substrate (BioRad #1705062) in ChemiDoc (BioRad). Densitometric analyses of the signals were carried out using ImageJ software.

### Inhibition of Glycosylation and In-vitro deglycosylation

J774A.1 macrophages were treated with 1µg/ml Tunicamycin (Sigma) for 3 or 12 hours following which the cells were harvested for immunoblotting. For in-vitro deglycosylation, 1 × 10^6^ cells were lysed and protein concentration was measured as described earlier. 20μg protein was denatured following manufacturer’s protocol followed by PNGase F and O-Glycosidase & Neuraminidase Bundle (NEB) treatment at 37°C for 1 hour. 4X NuPAGE loading buffer was added to the reaction mixture followed by immunoblotting as mentioned earlier.

### qRT–PCR of host Cu regulators

Total RNA was isolated from uninfected or *L. major*–infected macrophages using TRIzol reagent (Invitrogen #15596026). Verso cDNA synthesis kit (Thermo #AB1453A) was used for cDNA preparation from 1 μg of total RNA. All the primer were obtained from Eurofins and the details are listed in Table S1. Real-time PCR was performed with SYBR green fluorophore (Bio-Rad) using the 7500 real-time PCR system of Applied Biosystems. The relative transcript level of macrophage-specific genes was normalized using control set as the reference sample and GAPDH gene as an endogenous control. The experiments were performed as per minimum requirement of quantitative real-time PCR guidelines.

### Knockdown of ATP7A, CTR1 and ATOX1 with siRNA

Predesigned DsiRNA specific to mouse ATP7A (mm.Ri.Atp7a.13.1.), CTR1 (mm.Ri.Slc31a1.13.1), ATOX1 (mm.Ri.Atox1.13.1.) and Scrambled siRNA (Negative Control DsiRNA) from Integrated DNA Technologies were used for knockdown experiments. J774 macrophages were seeded on glass coverslip for amastigote nuclei counting and on 60 mm dishes for kDNA-based PCR method. Transfections were performed at 10 nM siRNA concentration using jetPRIME (Polyplus) transfecting reagent as per the manufacturer’s protocol and kept for 72 hours in serum free DMEM medium at 37°C in 5% CO2. Following this, the cells were washed twice with serum free DMEM medium and kept in DMEM with 10% FBS for another 12 hours. Knockdown of the candidates were confirmed by immunoblotting. Finally, they were infected *L. major* promastigotes for 12 hours and parasite load was calculated.

### Estimation of intracellular parasite burden

Infected macrophages were washed twice with 1X PBS and fixed with acetone:methanol (1:1). Coverslips were mounted on glass slides using 10µL of Fluoroshield with DAPI mountant to stain the nuclei of the fixed infected macrophages. Intracellular parasite burden represented as amastigotes/macrophage cell was quantified by counting the total number of DAPI-stained nuclei of macrophages and *L. major* amastigotes in a field (at least 100 macrophages were counted from triplicate experiments).

### kDNA-based parasite load estimation

Genomic DNA was isolated from the respective cell and tissue samples^55^. qRT-PCR was performed as mentioned earlier using *L. major* specific kDNA primers.

### Detection Cys-OH formed proteins by Cysteine Sulfenic Acid Probe

J774A.1 macrophages with or without infection (3 hour) were lysed in a specific lysis buffer containing DCP-Bio1 (Merck # NS1226) according to the manufacturer’s protocol. DCP-Bio1-bound proteins were pulled down with Streptavidin-coated magnetic beads (Genscript #L00936) overnight at 4 °C following manufacturer’s protocol. Biotinylated proteins were eluted in 1.5x NuPAGE LDS Sample buffer (Invitrogen #NP0007) containing 20 mM DDT (SRL #17315) and 2 mM biotin (SRL #18888) by heating at 95°C for 10 minutes and were determined by immunoblotting with CTR1 antibody.

### Manoeuvring copper bioavailability in mice and haemoglobin estimation

Six- to eight-week-old female BALB/c mice were divided into three groups. First group was Untreated (fed with regular diet and water), second group was fed with copper (20mg/L in drinking water) and third group was gavaged daily with the copper chelator, TTM (5mg/kg). For haemoglobin estimation, blood is withdrawn from retro-orbital plexus using capillary after 4 weeks and collected in EDTA solution from all three groups^56^. Hemocor-D (Coral Clinical systems) was used to measure the blood haemoglobin (Hb) level. According to the manufacturer’s protocol, 20 μl of blood is added to 5 ml of Hemocor-D solution and kept in room temperature for 5 mins followed by spectrophotometric reading at 540 nm.

### Mice infection and determination of lesion size and parasite load

The three groups were further divided into two categories, uninfected and infected. 1 × 10^6^ late log-phase *L. major* were suspended into PBS and injected into left hind footpad of infected groups. The progression of footpad lesion was monitored weekly post-infection by blindly measuring the left hind footpad with respect to the uninfected right hind footpad with a digital calliper^8^. At week 15 post-infection, mice were sacrificed, and parasite burden in the footpads of infected mice was quantified by kDNA-based PCR as mentioned earlier.

### Determination of cellular and tissue Cu concentration by ICP–MS

An equal number of J774A.1 macrophages were seeded in 60 mm culture dishes. After respective treatments, cells were washed with ice-cold PBS several six times and were harvested in centrifuge tubes. Following that, cells were counted, and an equal number of cells were digested in 100 μl of Nitric acid (Merck #1.00441.1000) overnight at 95°C. Copper standards were prepared from 23 Element standard (Reagecon #ICP23A20). Digested samples were diluted in 5 ml Milli Q water (Millipore) to bring the final concentration of nitric acid ≤2%, and analyzed using Quadrupole Inductively Coupled Plasma Mass Spectrometry (ICP-MS). For organ copper estimation, tissue was collected from euthanized mice following transcardial perfusion with PBS followed by flash-freezing in liquid nitrogen. 10 mg tissue for liver, heart, intestine, kidney, spleen and brain was digested and prepared for ICP-MS as mentioned above. For copper estimation from serum, 10 μl of serum was digested.

### Ceruloplasmin estimation

20 µl of the serum sample was pipetted into two different wells of two 96-well plates, each containing 80 µl of acetate buffer. For 5 mins, water bath at 30°C was used to submerge both plates marked as 5min and 15min respectively. O-dianisidine dihydrochloride reagent (Sisco Research Laboratories) was preincubated at 30°C and 25 µl was added into each well on the two plates and kept back in the waterbath. After exactly 5 min, the first plate was removed, and 230 µl of the 9 mol/L sulphuric acid (Merck Millipore) was added and mixed. The other plate was removed after exactly 15 minutes followed by addition of sulphuric acid. After that, spectrophotometric reading at 550nm was taken to detect the amount of oxidized *o-*dianisidine dihydrochloride.

### Image analysis

ImageJ, the image analysis software, (by Wayne Rasband) was used for analyzing the images^57^. Regions of interests were drawn manually on the best z-stack for each cell and the Colocalization_Finder plugin was used for colocalization studies. Pearson’s colocalization coefficient was measured using macro codes for quantifying colocalization^58^. Graphs were plotted using Graphpad Prism (version 9.4).

### Preparation of Illustrations

Illustrations were prepared in Inkscape 1.2.2. Liver - 201405 liver icon by Database Center for Life Science or DBCLS is licensed under CC BY 4.0 via Wikimedia Commons /modified. Heart - Heart-front icon by Servier https://smart.servier.com/ is licensed under CC-BY 3.0/modified. Proteasome-Proteasome by CristofferSevilla is licensed under CC BY 4.0/modified. Promastigote - Leishmania promastigote form by CristofferSevilla is licensed under CC BY 4.0 /modified. Amastigote - Leishmania amastigote form by CristofferSevilla is licensed under CC BY 4.0 /modified. Mice-Vector diagram of laboratory mouse (black and white) by Gwilz is licensed under CC BY-SA 4.0, via Wikimedia Commons /modified.

## Acknowledgement

This work was supported by DBT-Wellcome Trust India Alliance Fellowship (IA/I/16/1/502369) and Core Research Grant (CRG/2021/002150) from SERB, Department of Science and Technology (DST), Government of India and IISER K intramural funding to AG and Department of Biotechnology (DBT) and Department of Science and Technology (DST) grants BT/PR21170/MED/29/1109/2016 and EMR/2017/004506, respectively RD. CJC is a CIFAR fellow. CJC was supported by NIH grant GM79465. RP, SD and SM were supported by Pre-doctoral fellowship from Council of Scientific and Industrial Research and Raviranjan Panday by Pre-doctoral fellowship from Department of Biotechnology, India. SS was supported by Pre-doctoral fellowship from University Grants Commision, India. We thank IISER-Kolkata for providing us the facility to perform ICP-OES and IISER Pune for ICP-MS experiments. We thank IISER-Kolkata animal facility and Mr Anu Das for the mice experiments.

## Supplementary figures

**Figure S1.**
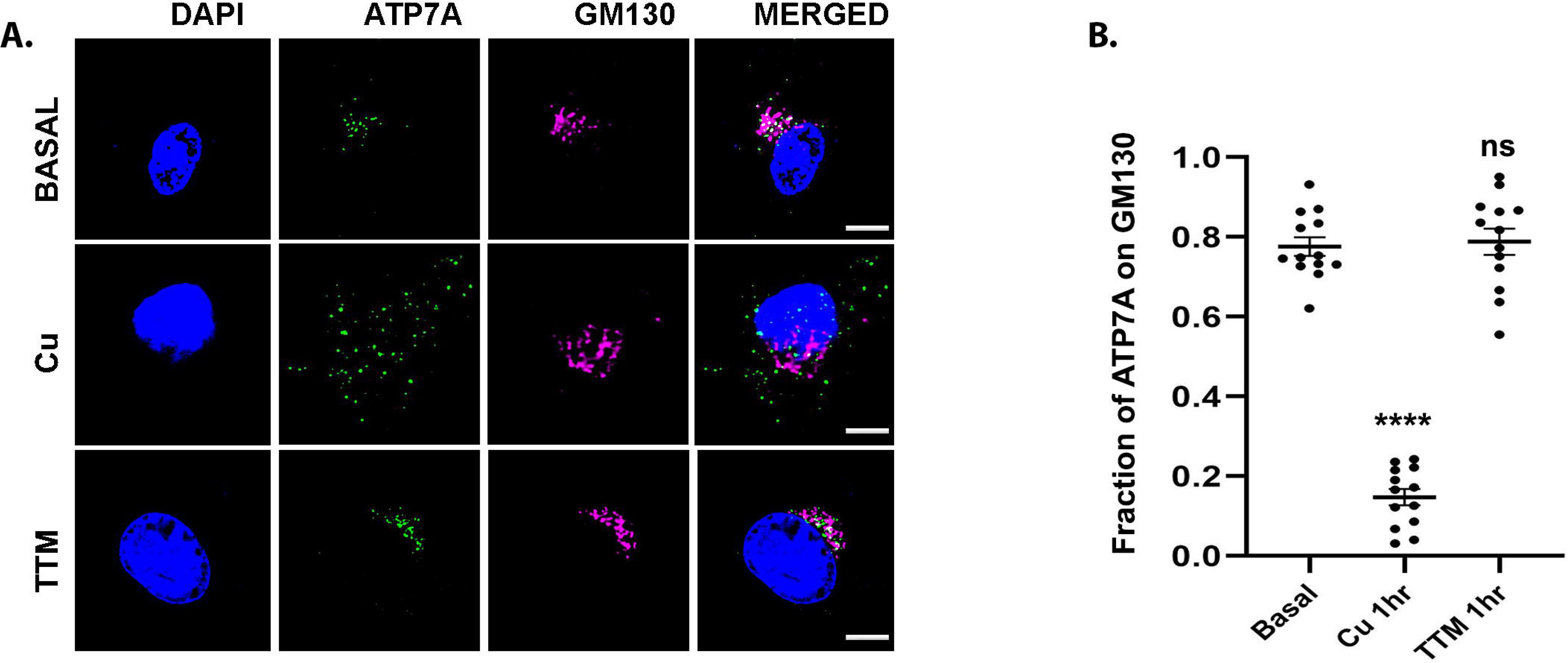
(A) Representative immunofluorescence image of ATP7A (green), co-stained with Golgi marker GM130 (magenta), in J774A.1 macrophages with no treatment, high copper (100 μM Cu) and copper chelated conditions (25 μM TTM) treatment for 2 hours. The merged images represent colocalization of ATP7A with GM130. The scale bar represents 5 μm. (B)) Fraction of ATP7A colocalization with GM130 from the above mentioned conditions is plotted. Sample size (n): 13. ****p ≤ 0.0001, ns; (Wilcoxon rank-sum test).

**Figure S2.**
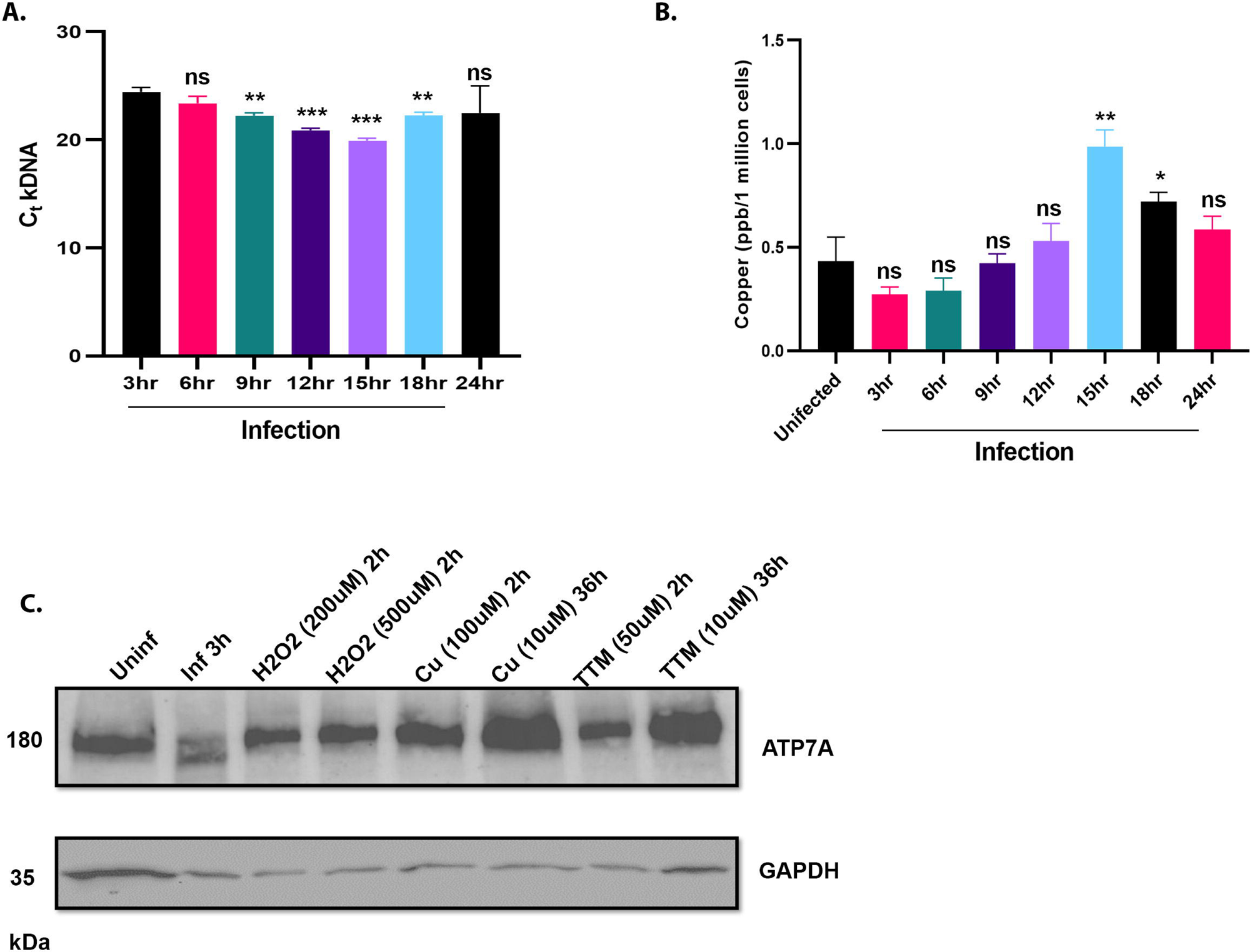
Determination of parasite load and copper content of macrophage upon *Leishmania* infection and *Leishmania*-specific modulation of host ATP7A. (A) *Leishmania* load was determined by C_t_ values of L.major kDNA at indicated time points after infection of J774A.1 macrophages and normalised against 3 hour infection. Error bars represent mean ± SD of values calculated from three independent experiments. **p ≤ 0.01, ***p ≤ 0.001, ns; (Student’s t test). (B) Measurement of intracellular copper level using ICP-MS at the above mentioned condtions. Error bars represent mean ± SD of values calculated from three independent experiments. *p ≤ 0.05, **p ≤ 0.01, ns; (Student’s t test). (C) Immunoblot of ATP7A of J774A.1 macrophages with and without 3 hour infection and indicated treatment conditions. GAPDH is used as housekeeping control.

**Figure S3.**
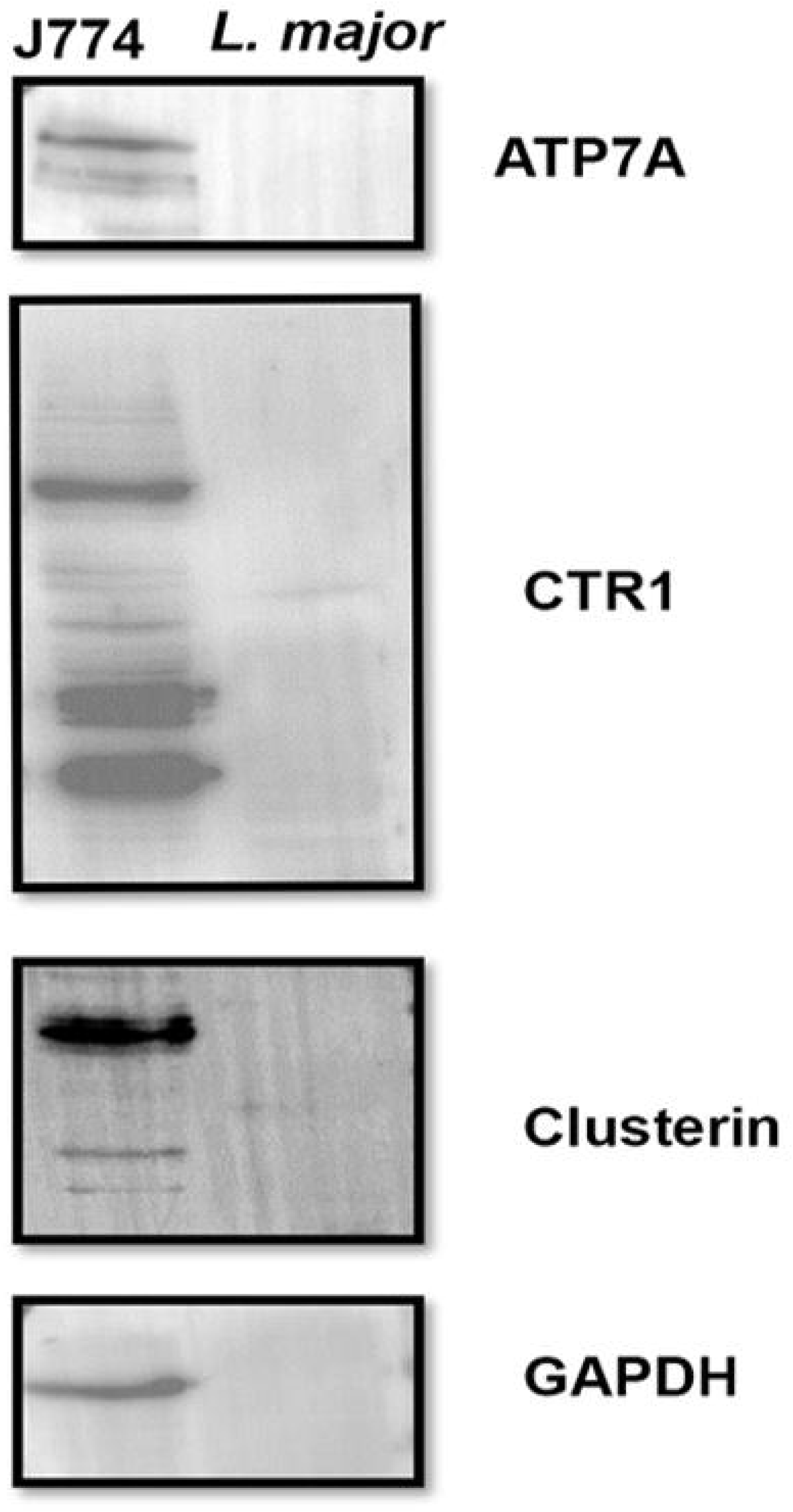
No crossreactivity of host specific antibodies with *Leishmania* proteins. Immunoblot of ATP7A, CTR1, Clusterin and GAPDH from J774A.1 and *L. major* probed using host specific antibodies, representing lack of crossreactivity of host specific antibodies with *Leishmania* proteins.

**Figure S4.**
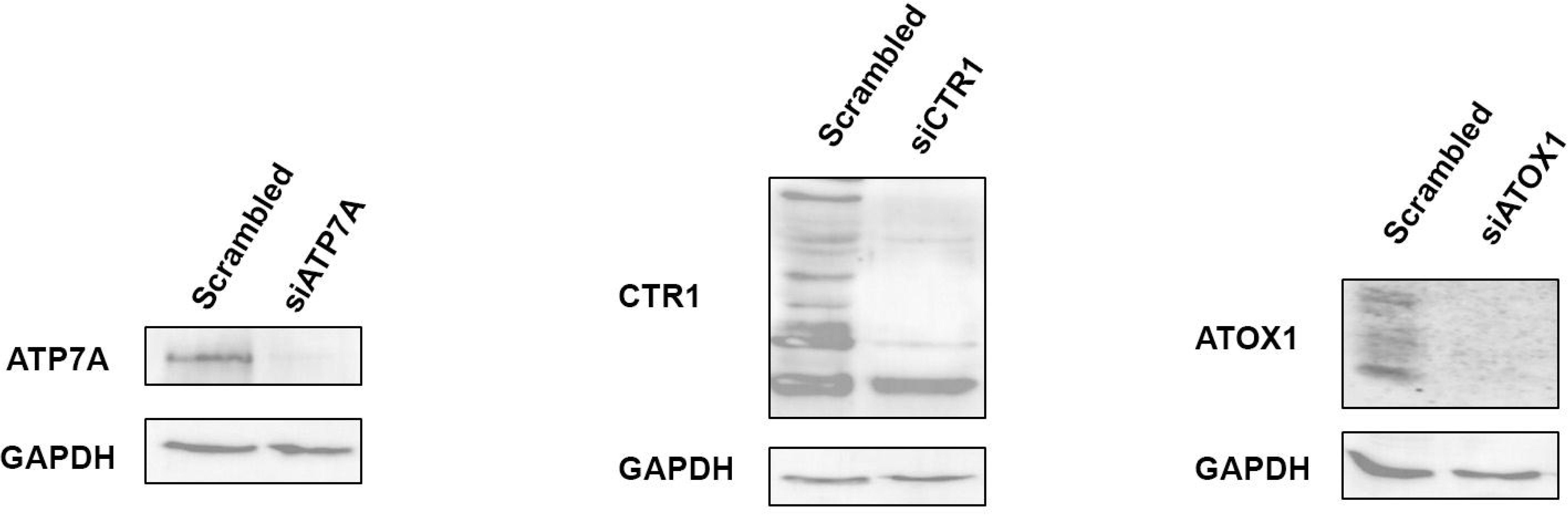
Confirmation of knockdown of host copper regulators via siRNA. Immunoblots of ATP7A, CTR1 and ATOX1 after treatment of J774A.1 macrophages with scrambled siRNA and ATP7A, CTR1 or ATOX1 siRNA respectively, confirming siRNA-mediated knockdowns. GAPDH is used as housekeeping control.

**Fig S5.**
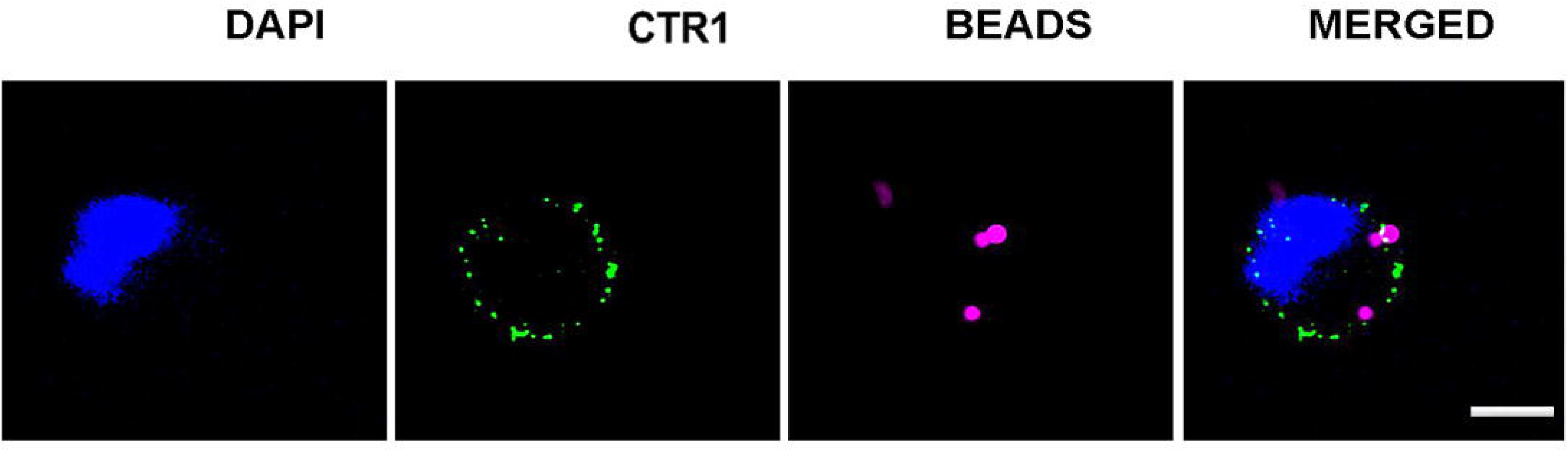
CTR1 endocytosis is not triggered by latex phagocytic beads. Immunofluorescence image of CTR1 (green) in J774A.1 macrophage with FluoSpheres™ Carboxylate-modified Microsphere beads (magenta) confirming absence of endocytosis of CTR1 due to general phagoscytosis triggered by the beads. Macrophage nuclei is stained with DAPI (blue).

**Fig S6.**
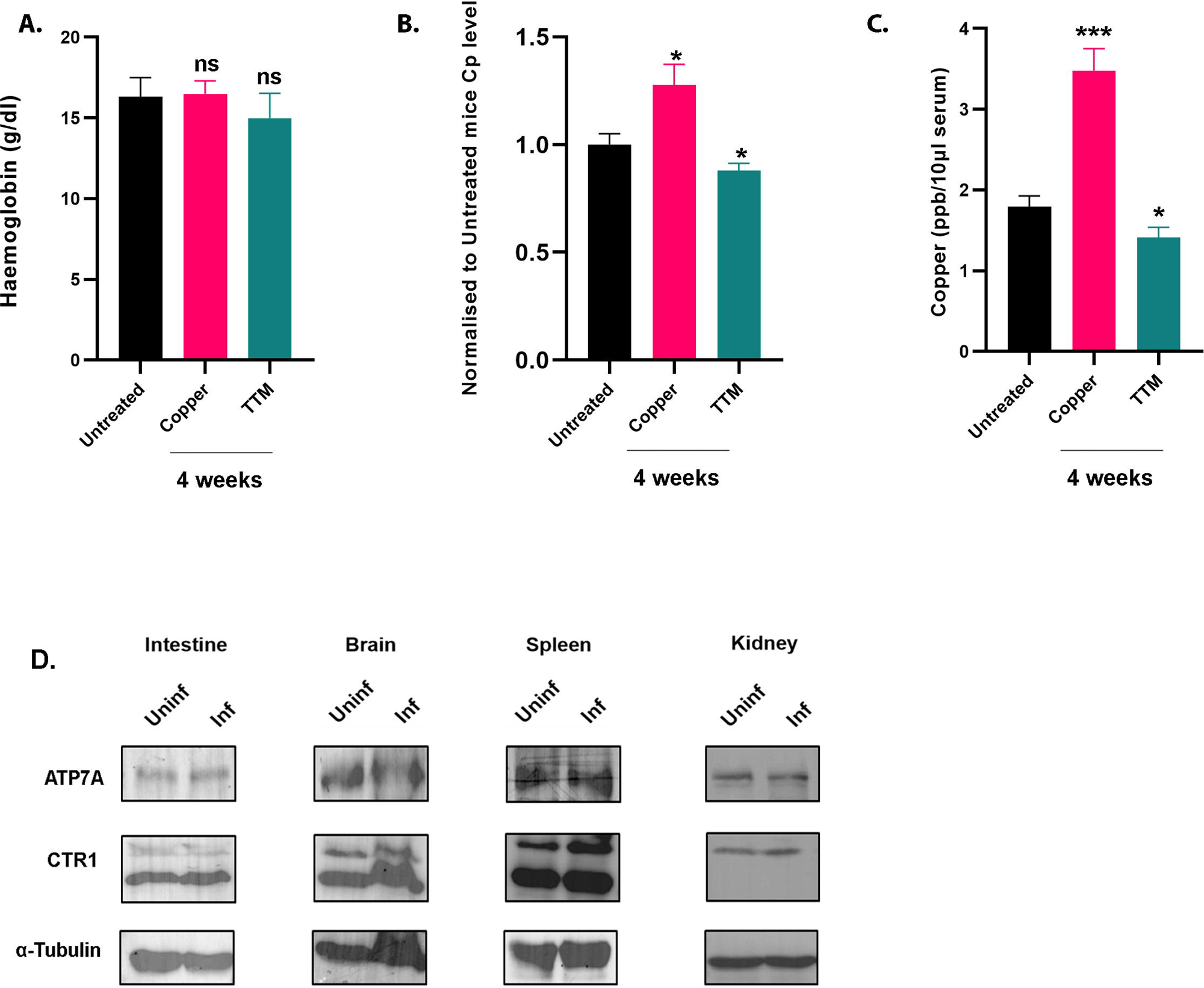
Optimal treatment of copper and TTM changes mice copper content without side effects, while *Leishmania* infection does not alter expression copper regulators in intestine, brain, spleen and kidney. Measurement of (A) Haemoglobin levels, (B) Ceruloplasmin level representative of serum bioavailable copper, normalised to that of the untreated mice and (C) whole serum copper of BALB/c mice 4 weeks post initiation of respective treatments to simulate different copper conditions. Error bars represent mean ± SD of values. Sample size (n): 3 for each condition, *p ≤ 0.05, ***p ≤ 0.001, ns; (Student’s t test). (D) Immunoblot of ATP7A and CTR1 from intestine, brain, spleen and kidney of infected and uninfected mice. The fold changes of ATP7A and CTR1 glycosylated monomer abundance normalized against housekeeping control α-tubulin have been mentioned.

## Notes

### Competing Interest Statement

The authors have declared no competing interest.

## References

1. Festa R.A. and Thiele D.J. Copper: An essential metal in biology. Curr. Biol. 2011;21:R877–R883. doi:10.1016/j.cub.2011.09.040.

2. Hodgkinson V. and Petris M.J. Copper homeostasis at the host-pathogen interface. J. Biol. Chem. 2012;287:13549–13555.

3. Nevitt T., Öhrvik H., Thiele D.J. Charting the travels of copper in eukaryotes from yeast to mammals. Biochim. Biophys. Acta. 2012;1823:1580–1593.

4. Kar S., Sen S., Maji S., Saraf D., Ruturaj, Paul R., Dutt S., Mondal B., Rodriguez-Boulan E., Schreiner R., Sengupta D., and Gupta A. 2022. Copper(II) import and reduction are dependent on His-Met clusters in the extracellular amino terminus of human copper transporter-1. Journal of Biological Chemistry. 298:101631.

5. Kuo Y.M., Zhou B., Cosco D., and Gitschier J. 2001. The copper transporter CTR1 provides an essential function in mammalian embryonic development. Proc. Natl. Acad. Sci. U.S.A. 98:6836– 6841.

6. Kaplan J.H., and Maryon E.B. 2016. How Mammalian Cells Acquire Copper: An Essential but Potentially Toxic Metal. Biophysical Journal. 110:7–13.

7. La Fontaine S., and Mercer J.F.B. 2007. Trafficking of the copper-ATPases, ATP7A and ATP7B: Role in copper homeostasis. Archives of Biochemistry and Biophysics. 463:149–167.

8. Paul R., Banerjee S., Sen S., Dubey P., Maji S., Bachhawat A.K., Datta R., and Gupta A. A novel leishmanial copper P-type ATPase plays a vital role in parasite infection and intracellular survival. J Biol Chem. 2022 Feb;298(2):101539.

9. Burza S., Croft S.L., and Boelaert M. Leishmaniasis. Lancet. 2018;392:951–970.

10. Okwor I. and Uzonna J. Social and economic burden of human leishmaniasis. Am. J. Trop. Med. Hyg. 2016;94:489–493.

11. Alvar J., Vélez I.D., Bern C., Herrero M., Desjeux P., Cano J., Jannin J., and den Boer M. Leishmaniasis worldwide and global estimates of its incidence. PLoS One. 2012;7

12. Peters N.C., Egen J.G., Secundino N., Debrabant A., Kimblin N., Kamhawi S., Lawyer P., Fay M.P., Germain R.N., and Sacks D. In vivo imaging reveals an essential role for neutrophils in leishmaniasis transmitted by sand flies. Science. 2008;321:970–974. Erratum in: Science 2008 Dec 12;322 (5908):1634.

13. Chang K.P. and Dwyer D.M. Leishmania donovani: Hamster macrophage interactions in vitro: Cell entry, intracellular survival, and multiplication of amastigotes. J. Exp. Med. 1978;147:515–530.

14. Haas A. The phagosome: Compartment with a license to kill. Traffic. 2007;8:311–330.

15. Thi E.P., Lambertz U., and Reiner N.E. Sleeping with the enemy: How intracellular pathogens cope with a macrophage lifestyle. PLoS Pathog. 2012;8

16. Gregory D.J., Sladek R., Olivier M., and Matlashewski G. Comparison of the effects of Leishmania major or Leishmania donovani infection on macrophage gene expression. Infect. Immun. 2008;76:1186–1192.

17. Beverley S.M. Hijacking the cell: Parasites in the driver’s seat. Cell. 1996;87:787–789.

18. Carrera L., Gazzinelli R.T., Badolato R., Hieny S., Muller W., Kuhn R., and Sacks D.L. Leishmania promastigotes selectively inhibit interleukin 12 induction in bone marrow-derived macrophages from susceptible and resistant mice. J. Exp. Med. 1996;183:515–526.

19. Croft S.L., Sundar S., and Fairlamb A.H. Drug resistance in leishmaniasis. Clin. Microbiol. Rev. 2006;19:111–126.

20. Sundar S. and Jaya J. Liposomal amphotericin B and leishmaniasis: Dose and response. J. Glob. Infect. Dis. 2010;2:159–166.

21. Festa R.A and Thiele D.J. (2012) Copper at the Front Line of the Host-Pathogen Battle. PLoS Pathog 8(9): e1002887.

22. Jones D.G., and Suttle N.F. Some effects of copper deficiency on leucocyte function in sheep and cattle. Res. Vet. Sci. 1981;31:151–156.

23. Crocker A., Lee C., Aboko-Cole G., and Durham C. Interaction of nutrition and infection: Effect of copper deficiency on resistance to Trypanosoma lewisi. J. Natl. Med. Assoc. 1992;84:697–706.

24. Newberne P.M., Hunt C.E., and Young V.R. The role of diet and the reticuloendothelial system in the response of rats to Salmonella typhilmurium infection. Br. J. Exp. Pathol. 1968;49:448–457.

25. Jones D.G., and Suttle N.F. The effect of copper deficiency on the resistance of mice to infection with Pasteurella haemolytica. J. Comp. Pathol. 1983;93:143–149.

26. Babu U. and Failla M.L. Copper status and function of neutrophils are reversibly depressed in marginally and severely copper-deficient rats. J. Nutr. 1990;120:1700–1709.

27. Babu U. and Failla M.L. Respiratory burst and candidacidal activity of peritoneal macrophages are impaired in copper-deficient rats. J. Nutr. 1990;120:1692–1699.

28. Wagner D., Maser J., Lai B., Cai Z., Barry C.E., 3rd, Höner Zu Bentrup K., and Russell D.G., Bermudez L.E. Elemental analysis of Mycobacterium avium-, Mycobacterium tuberculosis-, and Mycobacterium smegmatis-containing phagosomes indicates pathogen-induced microenvironments within the host cell’s endosomal system. J. Immunol. 2005;174:1491–1500.

29. White C., Lee J., Kambe T., Fritsche K., and Petris M.J. A role for the ATP7A copper-transporting ATPase in macrophage bactericidal activity. J. Biol. Chem. 2009;284:33949–33956.

30. Samanovic M.I., Ding C., Thiele D.J., and Darwin K.H. Copper in microbial pathogenesis: Meddling with the metal. Cell Host Microbe. 2012;11:106–115.

31. Argüello J.M., Mandal A.K., and Mana-Capelli S. Heavy metal transport CPx-ATPases from the thermophile Archaeoglobus fulgidus. Ann. N. Y. Acad. Sci. 2003;986:212–218.

32. Gupta A. and Lutsenko S. Evolution of copper transporting ATPases in eukaryotic organisms. Curr. Genomics. 2012;13:124–133.

33. Lutsenko S., Barnes N.L., Bartee M.Y., and Dmitriev O.Y. Function and regulation of human copper-transporting ATPases. Physiol. Rev. 2007;87:1011–1046.

34. Vanderwerf S.M., Cooper M.J., Stetsenko I.V. and Lutsenko S. Copper Specifically Regulates Intracellular Phosphorylation of the Wilson’s Disease Protein, a Human Copper-transporting ATPase. J. Biol. Chem; 2001. doi:10.1074/jbc.M102055200.

35. Petris M.J., Mercer J.F., Culvenor J.G., Lockhart P., Gleeson P.A. and Camakaris J. Ligand-regulated transport of the Menkes copper P-type ATPase efflux pump from the Golgi apparatus to the plasma membrane: a novel mechanism of regulated trafficking. EMBO J; 1996.Nov 15;15(22):6084–95.

36. Xiao T., Ackerman C.M., Carroll E.C, Jia S., Hoagland A., Chan J., Thai B., Liu C.S., Isacoff E.Y., and Chang C.J. Copper regulates rest-activity cycles through the locus coeruleus-norepinephrine system. Nat Chem Biol 14, 655–663 (2018). doi.org/10.1038/s41589-018-0062-z.

37. Manzl C., Enrich J., Ebner H., Dallinger R., and Krumschnabel G. Copper-induced formation of reactive oxygen species causes cell death and disruption of calcium homeostasis in trout hepatocytes. Toxicology. 2004 Mar 1;196(1-2):57–64. doi: 10.1016/j.tox.2003.11.001. PMID: 15036756.

38. Diaz-Albiter H., Sant’Anna M.R., Genta F.A., and Dillon R.J. Reactive oxygen species-mediated immunity against Leishmania mexicana and Serratia marcescens in the sand phlebotomine fly Lutzomyia longipalpis. J Biol Chem. 2012 Jul 6;287(28):23995–4003. doi: 10.1074/jbc.M112.376095. Epub 2012 May 29. PMID: 22645126; PMCID: PMC3390674.

39. Vonk W.I., de Bie P., Wichers C.G., van den Berghe P.V., van der Plaats R., Berger R., Wijmenga C., Klomp L.W., and van de Sluis B. The copper-transporting capacity of ATP7A mutants associated with Menkes disease is ameliorated by COMMD1 as a result of improved protein expression. Cell Mol Life Sci. 2012 Jan;69(1):149–63. doi: 10.1007/s00018-011-0743-1. Epub 2011 Jun 11. PMID: 21667063; PMCID: PMC3249196.

40. Materia S., Cater M.A., Klomp L.W., Mercer J.F., and La Fontaine S. Clusterin (apolipoprotein J), a molecular chaperone that facilitates degradation of the copper-ATPases ATP7A and ATP7B. J Biol Chem. 2011 Mar 25;286(12):10073–83.

41. Materia S., Cater M.A., Klomp L.W., Mercer J.F., and La Fontaine S. Clusterin and COMMD1 independently regulate degradation of the mammalian copper ATPases ATP7A and ATP7B. J Biol Chem. 2012 Jan 20;287(4):2485–99.

42. Kim B.E., Smith K., Meagher C.K., and Petris M.J. A conditional mutation affecting localization of the Menkes disease copper ATPase. Suppression by copper supplementation. J Biol Chem. 2002 Nov 15;277(46):44079–84. doi: 10.1074/jbc.M208737200. Epub 2002 Sep 6. PMID: 12221109.

43. Dawood A.A. and Altobje M.A. Inhibition of N-linked Glycosylation by Tunicamycin May Contribute to The Treatment of SARS-CoV-2. Microb Pathog. 2020 Dec;149:104586.

44. Suzuki T., Huang C., and Fujihira H. The cytoplasmic peptide:N-glycanase (NGLY1) - Structure, expression and cellular functions. Gene. 2016 Feb 10;577(1):1–7.

45. Lee H., Qi Y. and Im, W. Effects of N-glycosylation on protein conformation and dynamics: Protein Data Bank analysis and molecular dynamics simulation study. 2015 Sci Rep 5, 8926 (2015).

46. Inesi G. Calcium and copper transport ATPases: analogies and diversities in transduction and signaling mechanisms. J. Cell Commun. Signal. 2011. 5, 227–237.

47. Eisses J.F. and Kaplan J.H. Molecular characterization of hCTR1, the human copper uptake protein. J Biol Chem. 2002 Aug 9;277(32):29162–71. doi: 10.1074/jbc.M203652200. Epub 2002 May 28. PMID: 12034741.

48. Maryon E.B., Molloy S.A. and Kaplan J.H. O-linked glycosylation at threonine 27 protects the copper transporter hCTR1 from proteolytic cleavage in mammalian cells. J Biol Chem. 2007 Jul 13;282(28):20376–87. doi: 10.1074/jbc.M701806200. Epub 2007 May 24. PMID: 17525160.

49. Das A., Ash D., Fouda A.Y., Sudhahar V., Kim Y.M., Hou Y., Hudson F.Z., Stansfield B.K., Caldwell R.B., McMenamin M., Littlejohn R., Su H., Regan M.R., Merrill B.J., Poole L.B., Kaplan J.H., Fukai T., and Ushio-Fukai M. Cysteine oxidation of copper transporter CTR1 drives VEGFR2 signalling and angiogenesis. Nat Cell Biol. 2022 Jan;24(1):35–50. doi: 10.1038/s41556-021-00822-7. Epub 2022 Jan 13. PMID: 35027734; PMCID: PMC8851982.

50. Lee S., Howell S.B., and Opella S.J. NMR and mutagenesis of human copper transporter 1 (hCtr1) show that Cys-189 is required for correct folding and dimerization. Biochim Biophys Acta. 2007 Dec;1768(12):3127–34. doi: 10.1016/j.bbamem.2007.08.037. Epub 2007 Sep 21. PMID: 17959139; PMCID: PMC2275670.

51. Macintyre G., Gutfreund K.S., Martin W.R., Camicioli R., and Cox D.W. Value of an enzymatic assay for the determination of serum ceruloplasmin. J Lab Clin Med. 2004 Dec;144(6):294–301. doi: 10.1016/j.lab.2004.08.005. PMID: 15614251.

52. Stepien K.M. and Guy M. Caeruloplasmin oxidase activity: measurement in serum by use of o-dianisidine dihydrochloride on a microplate reader. Annals of Clinical Biochemistry. 2018; 55(1):149–157. doi:10.1177/0004563217695350

53. Pyatskowit J.W. and Prohaska J.R. Copper deficient rats and mice both develop anemia but only rats have lower plasma and brain iron levels. Comp Biochem Physiol C Toxicol Pharmacol. 2008 Apr;147(3):316–23. doi: 10.1016/j.cbpc.2007.11.008. Epub 2007 Dec 4. PMID: 18178529; PMCID: PMC2295218.

54. Kim B.E., Turski M.L., Nose Y., Casad M., Rockman H.A., Thiele D.J. Cardiac copper deficiency activates a systemic signaling mechanism that communicates with the copper acquisition and storage organs. Cell Metab. 2010 May 5;11(5):353–63. doi: 10.1016/j.cmet.2010.04.003. PMID: 20444417; PMCID: PMC2901851.

55. Myler P.J., Audleman L., deVos T., Hixson G., Kiser P., Lemley C., Magness C., Rickel E., Sisk E., Sunkin S., Swartzell S., Westlake T., Bastien P., Fu G., Ivens A., et al. Leishmania major Friedlin chromosome 1 has an unusual distribution of protein-coding genes. Proc. Natl. Acad. Sci. U. S. A. 1999;96:2902–2906.

56. Sharma A., Fish B.L., Moulder J.E., Medhora M., Baker J.E., Mader M., and Cohen E.P. Safety and blood sample volume and quality of a refined retro-orbital bleeding technique in rats using a lateral approach. Lab Anim (NY). 2014 Feb;43(2):63–6. doi: 10.1038/laban.432. PMID: 24451361; PMCID: PMC3989930.

57. Schneider C.A., Rasband W.S., Eliceiri K. NIH image to ImageJ: 25 years of image analysis. Nat. Methods. 2012;9:671–675.

58. McDonald J.H. and Dunn KW. Statistical tests for measures of colocalization in biological microscopy. J Microsc. 2013 Dec;252(3):295–302. doi: 10.1111/jmi.12093. Epub 2013 Oct 10. PMID: 24117417; PMCID: PMC4428547.

