## Supplemental Table 1 for "*Leishmania major*-induced alteration of host cellular and systemic copper homeostasis drives the fate of infection"

**Table S1. Oligonucleotides used in this paper**

| Primer | Primer Sequence (5’- 3’) |
| --- | --- |
| **ATP7AFP** | GAAGAGGTCGGACTGCTGTC |
| **ATP7ARP** | CCTTAGTAATGCCAACCTGAGAAGC |
| **CTR1FP** | CAAGATAGCCCGAGAGGGTC |
| **CTR1RP** | GATGTGCAGCACTGTCTGC |
| **COMMD1FP** | CAAGATAGCCCGAGAGGGTC |
| **COMMD1RP** | GATGTGCAGCACTGTCTGC |
| **ClusterinFP** | AGGAAAAGCCGTGCGGAAT |
| **ClusterinBP** | GCCTGGAGACATGTGGAGTT |
| **mGAPDHFP** | CGTGCCTGGAGAAACC |
| **mGAPDHRP** | TGGAAGAGTGGGAGTTGCTGTTG |
| **kDNA_FP** | AAGGGTGAACGCCAAAAACG |
| **kDNA_RP** | GTTCGGTTAATCCGCGAACG |
| **NGLY1FP** | ACTGTGCCAACAGGACCATC |
| **NGLY1RP** | GTGACTACGACACCCTAACCA |
