## Supplemental Table 2 for "*Leishmania major*-induced alteration of host cellular and systemic copper homeostasis drives the fate of infection"

| **Antibody Name** | **Type** | **Catalog Number** |
| --- | --- | --- |
| Rabbit Anti-ATP7A antibody | Primary | Abcam ab13995 |
| Rabbit Anti-CTR1 antibody | Primary | Abcam ab129067 |
| Rabbit Anti-COMMD1 antibody | Primary | Abcam ab102794 |
| Rabbit Anti Clusterin Beta Chain antibody | Primary | Abcam ab184099 |
| Rabbit Anti-GAPDH antibody | Primary | BioBharati BB-AB0060 |
| Mouse Anti-Lamp1 antibody | Primary | DSHB H4A3 |
| Mouse Anti-alpha 1 Sodium Potassium ATPase antibody | Primary | Abcam ab7671 |
| Rabbit Anti-Tubulin alpha antibody | Primary | Affinity Biosciences AF7010 |
| Anti-rabbit IgG, HRP-linked Antibody | Secondary | CST#7074 |
| Anti-mouse IgG, HRP-linked Antibody | Secondary | CST#7076 |
| Donkey Anti-Rabbit Alexa488 | Secondary | Life Technologies  A-21206 |
| Donkey Anti-Mouse Alexa568 | Secondary | Life Technologies  A-11057 |
